# Structural and functional insights into multiple BAM-bound conformations of BepA enabling substrate triage at the outer membrane

**DOI:** 10.64898/2026.08.05.742645

**Authors:** Ryoji Miyazaki, Mai Kimoto, Hidetaka Kohga, Takeru Nishi, Katsuhiro Sawasato, Yutaro S. Takahashi, Sho Asai, Yasushi Daimon, Takehiro Suzuki, Fina Amreta Laksmi, Yudhi Nugraha, Naoshi Dohmae, Hideki Shigematsu, Shin-ichiro Narita, Yoshinori Akiyama, Tomoya Tsukazaki

**Affiliations:** Nara Institute of Science and Technology, Ikoma, Nara, Japan; Department of Life Sciences, Faculty of Agriculture, Iwate University, Morioka, Iwate 020-8550, Japan; Institute for Life and Medical Sciences, Kyoto University, Kyoto 606-8507, Japan; Biomolecular Characterization Unit, Technology Platform Division, RIKEN Center for Sustainable Resource Science, Wako, Saitama 351-0198, Japan; Directorate of Laboratory Management, Research Facilities, and Science Technology Parks, National Research and Innovation Agency (BRIN), Cibinong, Bogor, West Java, Indonesia; Research Center for Molecular Biology Eijkman, National Research and Innovation Agency (BRIN), Cibinong, Bogor, West Java, Indonesia; Structural Biology Division, Japan Synchrotron Radiation Research Institute, Sayo, Hyogo, Japan; Faculty of Health and Nutrition, Yamagata Prefectural Yonezawa University of Nutrition Sciences, 992-0025 Yamagata, Japan

**Author notes:** Correspondence (T.T.), (R.M.), (Y.A.). equally contributed (Mai Kimoto, Hidetaka Kohga).

## Abstract

The outer membrane (OM) of Gram-negative bacteria acts as a selective permeability barrier against toxic compounds, and for this function, requires proper assembly of outer membrane proteins (OMPs) by the β-barrel assembly machinery (BAM) complex. A periplasmic metalloprotease BepA promotes maturation of LptD, an essential OMP, while degrades LptD intermediates aberrantly stalled at BAM. However, how BepA switches between these functions remains unclear. Here, we report cryo-EM structures of BAM– BepA complexes that capture multiple BepA conformations. *In vivo* crosslinking and cysteine-accessibility analyses show that possible regulatory elements, α6- and α9-loops that cover the proteolytic active site, can assume open conformations in living cells. Functional analyses reveal that α6-loop opening supports substrate interaction, degradation, and membrane association, whereas α9-loop opening is required for proteolytic activation and stabilizes BAM association. These findings provide insights into how distinct loop rearrangements regulate BepA-mediated substrate triage at BAM.

## Introduction

The cell envelope of Gram-negative bacteria consists of two biological membranes, the inner membrane (IM) and the outer membrane (OM), and the periplasmic space between them, which contains the peptidoglycan layer. The OM is directly exposed to the external environment and functions as an essential selective permeability barrier against toxic compounds, including antibiotics (*1*). Its integrity depends on the coordinated biogenesis of β-barrel outer membrane proteins (OMPs) and lipopolysaccharide (LPS), two major constituents of the OM (*2*). OMP assembly and LPS transport to the OM are mediated by two essential envelope machineries, the β-barrel assembly machinery (BAM) complex and the LPS transport (Lpt) system, respectively.

Following synthesis in the cytoplasm, nascent OMPs are translocated across the IM by the Sec machinery, and their signal sequences are cleaved off by signal peptidase during or just after membrane translocation (*3–6*). Periplasmic chaperones, including SurA and Skp, subsequently deliver unfolded OMPs to the BAM complex at the OM (*7*, *8*). In *Escherichia coli* (*E. coli*), the BAM complex consists of two essential components, the OMP BamA and the lipoprotein BamD, and three nonessential OM lipoproteins, BamB, BamC, and BamE (*9*, *10*). BamA contains five flexible N-terminal polypeptide transport-associated (POTRA) domains and a C-terminal membrane-embedded β-barrel domain with a lateral gate that can open and close for OMP assembly. Structural and biochemical studies have revealed key steps in BAM-mediated OMP biogenesis. SurA initially delivers unfolded OMPs into the BAM complex by interacting with the POTRA1 domain of BamA, BamB, and BamE (*11–13*). BamD has been proposed to initially recognize internal signals within substrate OMPs and organize their C-terminal β-strands for subsequent engagement with the lateral gate of BamA (*14*). The C-terminal β-signal is then recognized at the lateral gate of BamA, where the substrate folds into a β-barrel and is subsequently released laterally into the OM (*15–19*).

LPS is synthesized at the inner leaflet of the IM, flipped to the periplasmic side by MsbA, and transported from the IM to the OM by the Lpt proteins (LptA, LptB, LptC, LptF, and LptG) (*20*, *21*). At the OM, LptD, an essential β-barrel OMP, forms the LPS translocon together with the lipoprotein LptE and plays an essential role in facilitating LPS transport to the outer leaflet of the OM (*22*, *23*). LptD has two forms: a mature form (LptD^NC^) and an intermediate form (LptD^C^), which contain two intramolecular disulfide bonds between nonconsecutive (NC) and consecutive (C) Cys pairs, respectively (*22*, *24*). During LptD maturation, LptD^C^ is first generated in the periplasm and is subsequently converted to LptD^NC^ during its assembly mediated by the BAM complex (*24–26*). In addition to LptE, LptM, a recently identified LptD-interacting lipoprotein, is involved in this conversion process (*27*, *28*). LptD maturation is a complex process that can generate stalled or aberrant assembly intermediates. Several proteases act at distinct stages of LptD quality control, including DegP in the periplasm and BepA and YcaL at the BAM complex (*29*).

BepA is a periplasmic metalloprotease that comprises an N-terminal protease domain and a C-terminal tetratricopeptide repeat (TPR) domain, and plays a crucial role in maintaining OM integrity (*24*). Genetic and biochemical studies have suggested that BepA is a bifunctional quality-control factor having protease and chaperone-like activities that act during LptD maturation (*24*). At the BAM complex, BepA associates with an LptD assembly intermediate via its edge-strand and either promotes its maturation in a chaperone-like manner under normal conditions or degrades it when its assembly is aberrantly stalled (**Fig. 1A**) (*24*, *30*, *31*). However, the molecular mechanism of substrate triage by BepA remains unclear. The crystal structure of BepA has provided important insights into its regulatory mechanism. In the BepA crystal structure, the proteolytic catalytic site and the substrate-recognition edge-strand are covered by two loop regions, α6- and α9-loops (**Fig. 1B and C**) (*32*), suggesting that these loops are key regulatory elements of BepA function. Indeed, the conserved His-246 residue in the α9-loop is coordinated to the catalytic Zn ion of BepA and inhibits its protease activity, probably enabling BepA to act as a chaperone-like factor without degrading substrates (His-switch) (*33*, *34*). Thus, conformational rearrangements of the α6- and α9-loops are likely central to the switch between the chaperone-like and protease functions of BepA. However, the functional role of the α6-loop and how α6- and α9-loop movements are regulated remain unclear.

**Fig. 1.**
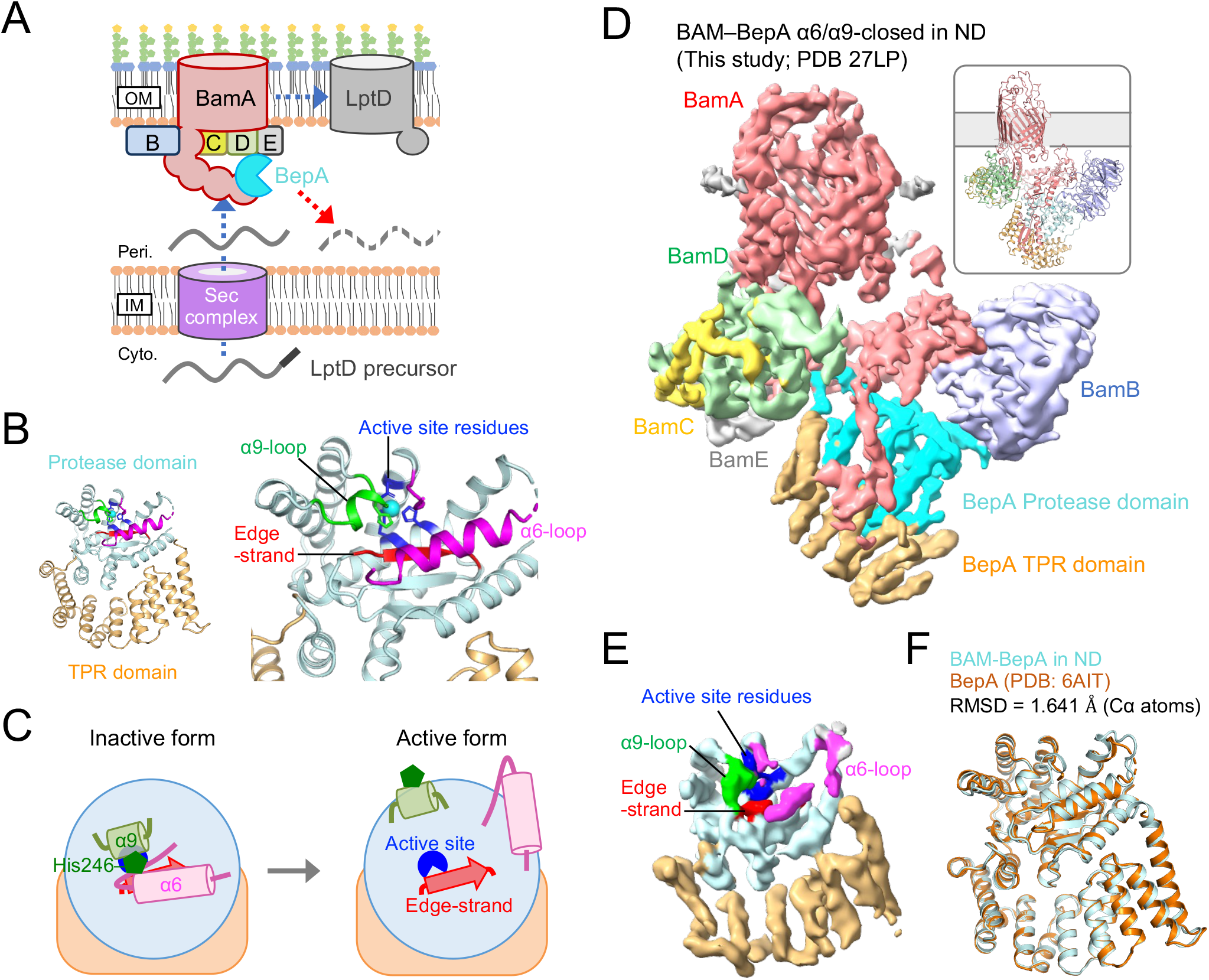
Cryo-EM structure of the nanodisc-reconstituted BAM–BepA complex. (**A**) Schematic model for BepA-mediated quality control of an LptD assembly intermediate. (**B**) Crystal structure of BepA (PDB code: 6AIT). The protease and the TPR domains of BepA are shown in light cyan and orange, respectively. The edge-strand, the proteolytic active-site residues (the HExxH motif and the third zinc ligand, Glu-201), the α6-loop, and the α9-loop in the protease domain are shown in red, blue, magenta, and green, respectively, and the coordinated zinc atom is shown in cyan. (**C**) Schematic model of BepA activation through α6- and α9-loop opening. (**D**) Cryo-EM structure of BAM–BepA complex. A Cryo-EM map and a cartoon model of the nanodisc-reconstituted BAM–BepA complexes are shown (PDB: 27LP). BamA, BamB, BamC, BamD, and BamE are shown in red, light blue, yellow, light green, and gray, respectively. The BepA protease and TPR domains are shown in cyan and orange, respectively. (**E**) Cryo-EM map of BepA in the BAM–BepA complex shown in (D). The BepA regions are colored as in (B). (**F**) Structural comparison of BepA in the nanodisc-reconstituted BAM–BepA complex with the BepA crystal structure. BepA in the BAM–BepA complex and the BepA crystal structure are shown in light cyan and orange, respectively.

Here, we investigated the molecular mechanism underlying BepA function and the associated conformational changes at the BAM complex. Cryo-EM and biochemical analyses revealed multiple BAM–BepA conformations, including the α6/α9-closed, α6-open/α9-closed, and α6/α9-open states. *In vivo* photo-crosslinking and accessibility analyses suggested that BepA can adopt the α6-open state at the BAM complex in living cells, while the α9-loop remains closed until a later activation step. Functional analyses further show that α6-loop opening is required for substrate interaction, substrate degradation, and membrane-association of BepA, whereas α9-loop opening, which is essential for activation of its proteolytic function, further stabilizes BepA association with the membrane. These results suggest that BepA regulates substrate triage at the BAM complex through separable conformational changes of the α6- and α9-loops.

## Results

### BepA adopts a closed conformation in the nanodisc-reconstituted BAM–BepA complex

To elucidate the molecular basis of the interaction between BepA and the BAM complex, we determined the structure of the BAM–BepA complex. Previous photo-crosslinking analyses indicated that the TPR domain of BepA contacts BamA, BamC, BamD, and that the POTRA domain of BamA contacts BepA (*30*). Thus, to stabilize the BAM–BepA interaction by disulfide crosslinking, we introduced cysteine substitutions at BepA F404 in the TPR domain and BamA P47 in the POTRA1 domain (**fig. S1A**). We observed that BepA F404C degraded overproduced LptD (**fig. S2A**) and BamA P47C rescued the conditional lethal BemA-depleted strain (**fig. S2B**). Thus, these cysteine substitutions did not affect the functions of BepA or BamA (**fig. S2A and B**). We also introduced the catalytically inactive E137Q mutation into BepA because this mutation stabilizes the interaction of BepA with BAM (*24*, *30*). We coexpressed BamA(P47C), BamB, BamC, BamD, BamE, and BepA(F404C, E137Q) carrying a His_10_-tag at the N-terminus of the mature protein, purified the resulting BAM–BepA complex, reconstituted it into lipid nanodiscs using a membrane scaffold protein (MSP1D1), and determined its structure by cryo-EM single-particle analysis (**Fig. 1D**; **figs. S1A–G**). In this structure, the TPR domain of BepA was located at the periplasmic side, whereas the protease domain was positioned toward the OM side of the complex. The α6- and α9-loops folded over the BepA catalytic site, consistent with the closed conformation previously observed in the crystal structure of BepA (**Fig. 1E and F**) (*32*). Notably, despite its association with the BAM complex, BepA retained this closed conformation, suggesting that association with BAM alone is insufficient to induce the active conformation of BepA.

The C-terminally fused His_10_-tag of BepA is self-cleaved *in vivo*(*24*, *30*, *31*), suggesting that the C-terminal tag may act as a substrate-like element and influence the movements of the α6- and α9-loops required for BepA to adopt its active conformation. Therefore, we performed cryo-EM analysis of the BAM–BepA complex containing BepA carrying both an N-terminal His_10_ tag and an additional C-terminal His_10_ tag. However, the nanodisc-reconstituted BAM–BepA structures are essentially identical in the presence and absence of the C-terminal His₁₀ tag on BepA, indicating that the C-terminal tag does not markedly alter the closed conformation of BepA (**fig. S3**). Thus, the nanodisc-reconstituted BAM–BepA structure captured a BAM-bound yet catalytically inactive state of BepA. This raised the possibility that BepA adopts additional conformations during its functional interactions with BAM *in vivo*.

### BepA adopts additional BAM-bound conformations *in vivo*

Our structural analyses described above imply that BepA needs to undergo conformational changes to exert its functions. To get insight into possible structural changes of BepA in living cells, we performed *in vivo* photo-crosslinking analysis. We arbitrarily introduced *p*-benzoyl-L-phenylalanine (*p*BPA) in the protease domain of BepA(E137Q)-His_10_ and expressed BepA(*p*BPA)-His_10_ derivatives in Δ*bepA* cells. After UV-irradiattion, total cellular proteins were analyzed by SDS-PAGE and immunoblotting with anti-BepA antibodies. We identified that 16 BepA(*p*BPA) derivatives generated crosslinked products with higher apparent molecular masses than that of uncrosslinked BepA-His_10_ (**Fig. 2A**). Immunoblotting with anti-BamA antibodies indicated that several residues in the BepA protease domain were close to BamA, as previously observed for BepA F404*p*BPA in the TPR domain (**Fig. 2a**)(*30*). Immunoblotting with anti-BamB antibodies showed that BepA V162*p*BPA and R214*p*BPA were crosslinked to BamB (**fig. S4A**). In addition, immunoblotting with anti-BamD antibodies showed that BepA S63*p*BPA was crosslinked to BamD (**fig. S4B**). These BepA(*p*BPA) derivatives suppressed the accumulation of LptD^C^, an LptD assembly intermediate, in Δ*bepA* cells when expressed from a plasmid, indicating that they were functional and likely reflected the physiological behavior of BepA (**fig. S4C**).

**Fig. 2.**
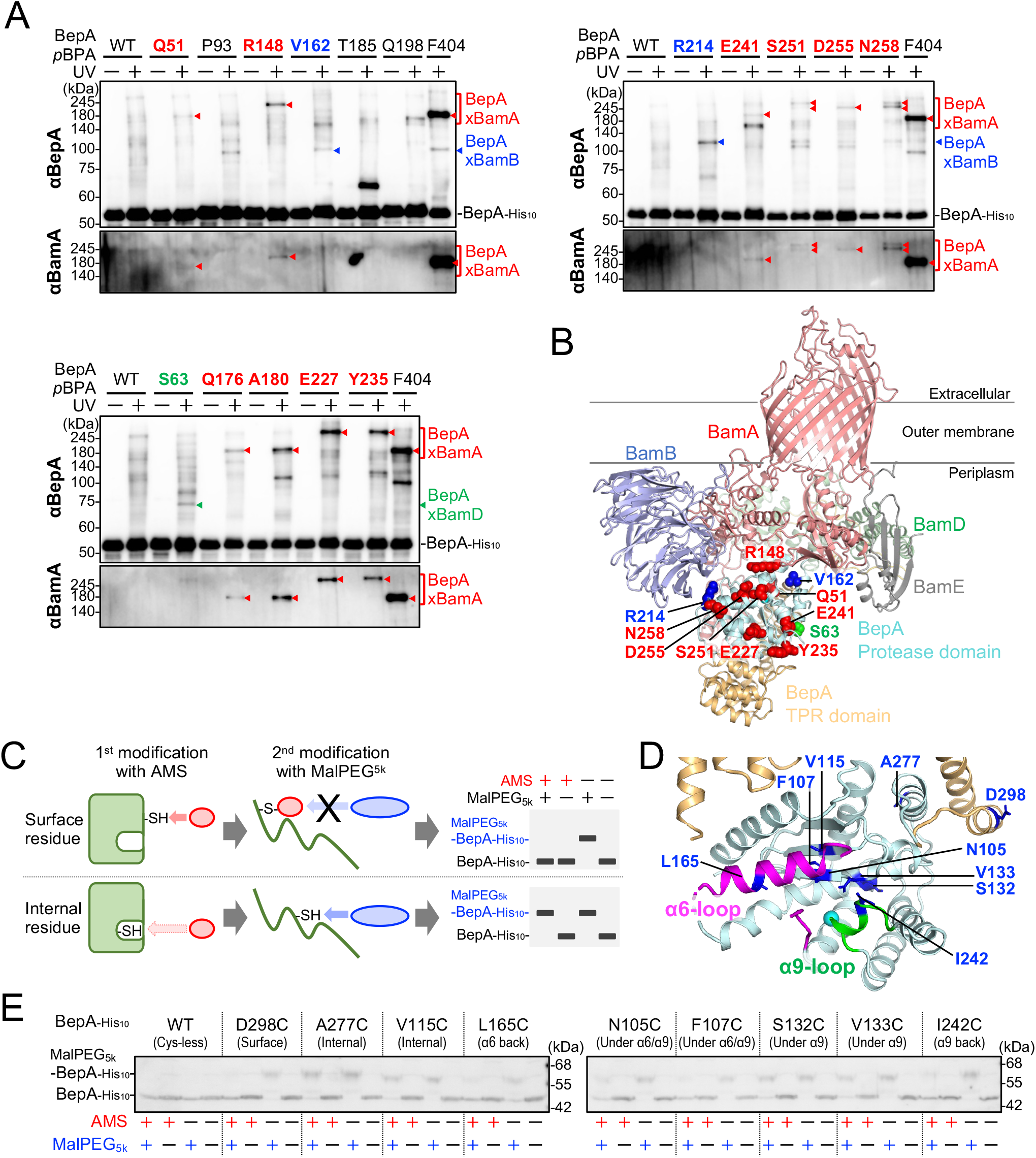
In vivo analyses suggest additional BAM-bound conformations of BepA. (**A**) *In vivo* photo-crosslinking analysis of BepA. Cells of SN56 (Δ*bepA*) carrying both pEVOL-pBpF and pUC18-*bepA(amb, E137Q)-his_10_* were grown at 30 °C in LB medium containing 0.5 mM *p*BPA until early log phase and induced with 1 mM IPTG for 1 h to express the indicated BepA(*p*BPA) variants. Cultures were then divided into two portions treated with or without UV irradiation for 10 min at 4 °C. Total cellular proteins were acid-precipitated, solubilized with SDS buffer, and subjected to pull-down with Ni-NTA agarose. The purified proteins were analyzed using SDS–PAGE followed by immunoblotting using the indicated antibodies. (**B**) Mapping of the crosslinking sites on the nanodisc-reconstituted BAM–BepA structure. BepA residues crosslinked with BamA, BamB, and BamD are indicated by red, blue, and green spheres, respectively. (**C**) Schematic representation of the AMS/MalPEG modification assay. (**D**) Cysteine substitution sites in BepA. In the crystal structure of BepA (PDB: 6AIT), the α6- and α9-loops are shown in magenta and green, respectively. The introduced cysteine positions are shown as blue sticks. (**E**) AMS/MalPEG modification assay using BepA Cys derivatives. Cells of SN56 (Δ*bepA*) carrying pHSG575-*bepA(mut)-his_10_* were grown at 30 °C in LB medium until early log phase, induced with 1 mM IPTG for 2 h to express the indicated BepA variants, converted to spheroplasts and treated with AMS. After quenching the AMS, proteins were acid-precipitated, denatured in SDS, and further treated with MalPEG_5k_ to modify free thiols. The samples were then analyzed by 7.5% Laemmli SDS-PAGE and immunoblotting with anti-BepA antibodies.

We then mapped the identified BamA-, BamB-, and BamD-crosslinking sites onto the nanodisc-reconstituted BAM–BepA structure. BepA R214 was indeed located near BamB, consistent with the observed photo-crosslinking. However, several other crosslinking residues were distant from their identified crosslinking partners in the nanodisc-reconstituted BAM–BepA structure (**Fig. 2B**). These results indicate that BepA can also adopt additional BAM-bound conformations alongside the nanodisc-reconstituted structure in living cells.

### The α6-loop undergoes dynamic conformational changes *in vivo*

Although the α6- and α9-loops cover the BepA catalytic site in the nanodisc-reconstituted BAM– BepA structure, BepA-mediated proteolysis requires these loops to move away from the catalytic site. Therefore, we performed an AMS accessibility assay to examine whether the α6- and α9-loops undergo conformational changes in living cells. In this assay, a cysteine substitution was introduced at a position of interest in BepA. The protein was first treated with a small thiol-modifying reagent, such as AMS, under native conditions. After denaturation, the protein was further reacted with methoxypolyethylene glycol 5000 maleimide (MalPEG_5k_), a bulky 5-kDa thiol-modifying reagent, and then analyzed by SDS-PAGE and immunoblotting. If the introduced cysteine is exposed, it should be accessible to modification by AMS and therefore unavailable for MalPEG modification. In contrast, if the introduced cysteine is buried inside the protein, it should remain inaccessible to AMS under native conditions and become available for modification by MalPEG after denaturation, resulting in an upward band shift on SDS-PAGE (**Fig. 2C**).

We introduced a cysteine substitution at several positions in BepA, including residues covered by α6- and α9-loops (**Fig. 2D**). As controls, cysteine introduced at the surface-exposed residue D298 was modified by AMS, whereas cysteines introduced at the buried residues A277 and V115 were not modified by AMS under native conditions and were subsequently modified by MalPEG_5k_ after denaturation (**Fig. 2E**). These results confirmed that the assay can distinguish exposed and buried residues in BepA. We next examined the accessibility of residues that are covered by the α6- and α9-loops. Cysteines introduced at S132 and V133, which are buried under the α9-loop in the closed structure, were only weakly modified by AMS. In contrast, cysteines introduced at the other positions, including those around the α6-loop-covered region, were efficiently modified by AMS (**Fig. 2E**). This weak modification was not because S132C and V133C were intrinsically unreactive, but rather because their accessibility was limited by the α9-loop, as their AMS modification markedly increased in the α9-deletion variant (**fig. S5**). All BepA cysteine variants retained protease activity comparable to that of wild-type BepA (**fig. S2C**), suggesting that their behavior reflects the physiological conformational dynamics of BepA. These results suggest that, in living cells, the α9-loop remains relatively closed, whereas the region covered by the α6-loop becomes accessible, consistent with an open or dynamic conformation. Together, the *in vivo* photo-crosslinking and AMS-accessibility analyses suggest that BepA adopts BAM-bound conformations distinct from the nanodisc-reconstituted closed state.

### Cryo-EM structures capture α6-open and α6-closed BAM–BepA conformations

To structurally capture the additional BAM-bound conformations of BepA suggested by the *in vivo* analyses, we next analyzed the BAM–BepA complex under DDM-solubilized conditions. In the nanodisc-reconstituted structure, BepA is positioned relatively peripherally on the periplasmic side of BAM (**Fig. 1C**). However, the photo-crosslinking results suggested that BepA can also associate more deeply with the BAM complex (**Fig. 2A and B**). We considered that the MSP1D1 in nanodiscs might restrict conformational rearrangements around the periplasmic region of BAM. Therefore, we purified the same BAM–BepA complexes, subjected them to cryo-EM analysis in DDM-solubilized conditions (**fig. S6**), and determined two BAM–BepA structures that were distinct from the nanodisc-reconstituted BAM–BepA structure (**Fig. 3A and B**). In one structure, the α6-loop remained closed, and BepA was positioned more deeply within the BAM complex than in the nanodisc-reconstituted structure. In the other structure, the α6-loop region adopted an open conformation and extended toward the BamA β-barrel, with BepA inserted further into the periplasmic region of the BAM complex. We refer to these structures as the α6-closed and α6-open BAM–BepA structures, respectively. Among the three BAM–BepA structures determined in this study, the α6-open structure was more consistent with the *in vivo* photo-crosslinking results. Most crosslinked residues were located close to their respective crosslinking partners among the Bam subunits (**Fig. 3C**). In this structure, the protease domain of BepA, including its N-terminal region, contacts the POTRA2–5 domains of BamA (**fig. S7**). In particular, the opened α6-loop region was positioned near the BamA β-barrel in this structure, consistent with the observation that residues within the α6-loop were crosslinked to BamA (**Fig. 2A and 3C**; **fig. S7**). In addition, the TPR domain of BepA was positioned close to BamA POTRA1 and POTRA2, as well as BamC and BamD, consistent with our previous *in vivo* photo-crosslinking analysis targeting the TPR domain of BepA (**fig. S8**) (*30*). Taken together, these results suggest that the α6-open BAM–BepA structure represents one physiologically relevant BAM-bound conformation of BepA.

**Fig. 3.**
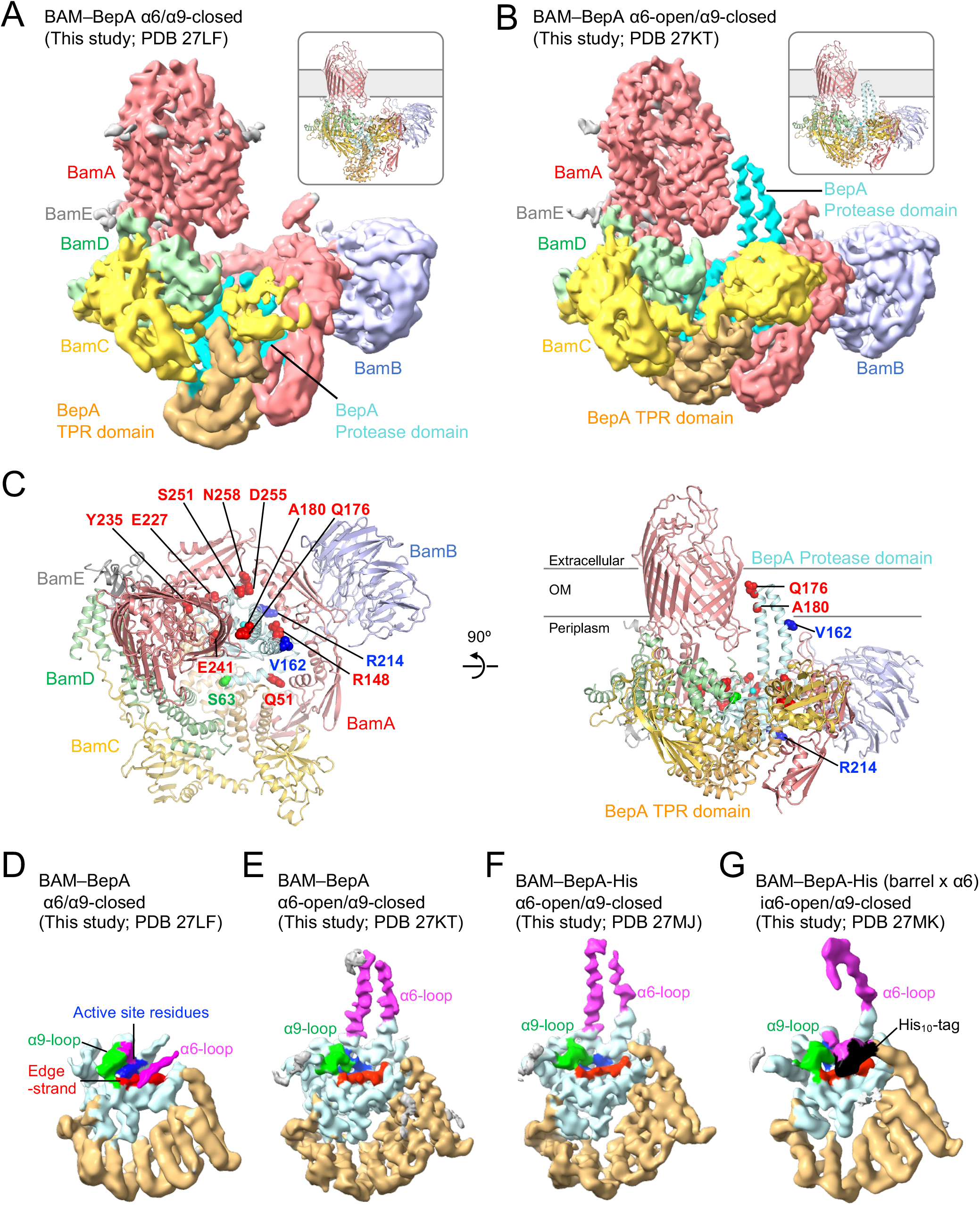
Cryo-EM structures of DDM-solubilized BAM–BepA reveal α6-closed and α6-open conformations. (**A, B**) Cryo-EM maps and cartoon models of DDM-solubilized BAM–BepA complexes. (A) BAM–BepA α6/α9-closed structure (PDB: 27LF). (B) BAM–BepA α6-open/α9-closed structure (PDB: 27KT). BamA, BamB, BamC, BamD, BamE, and the protease and TPR domains of BepA are colored as in Fig. 1C. (**C**) Mapping of the crosslinking sites on the BAM–BepA α6-open/α9-closed structure. BepA residues crosslinked with BamA, BamB, and BamD are indicated by red, blue, and green spheres, respectively. (**D– G**) Cryo-EM maps of BepA in BAM–BepA complexes. (D) BAM–BepA α6/α9-closed structure. (E) BAM–BepA α6-open/α9-closed structure. (F) BAM–BepA-His α6-open/α9-closed structure. (G) β-barrel-α6-crosslinked BAM–BepA-His α6-open/α9-closed structure. The BepA regions are colored as in Fig. 1b. The cryo-EM map corresponding to the C-terminal His_10_-tag is shown in black in (G).

In both the α6-closed and α6-open structures, the α9-loop continued to cover the catalytic site (**Fig. 3D and E**). This observation was consistent with the AMS accessibility assay (**Fig. 2E**), which suggested that the α9-loop remains relatively closed in living cells. To examine whether a substrate can access the catalytic site in this state, we also performed structural analysis using BepA with a C-terminal His_10_-tag, which can behave as a substrate-like element (*24*, *30*, *31*). A similar α6-open/α9-closed structure was obtained (**fig. S9**). However, no additional cryo-EM map corresponding to the C-terminal His_10_-tag was observed near the catalytic site (**Fig. 3F**). In a previously reported AlphaFold model model of BAM– BepA(*35*), the α6-loop of BepA is also open, but interacts with a different site on the BamA β-barrel (**Fig. 3B**; **fig. 10A**). Our BamA-BepA disulfide-crosslinking analysis demonstrated that BepA interacts with BamA as predicted by this model (**fig. 10B**). We stabilized the BamA–BepA-His_10_ interaction by disulfide crosslinking between BamA P779C and BepA A180C and performed cryo-EM analysis to obtain structural information on another α6-loop open conformation (**fig. S11**). In this structure (**Fig. 3G**), the α6-loop region was indeed open, although it adopted a slightly different conformation from that in the α6-open BAM– BepA structures (**Fig. 3F and G**). Notably, an additional cryo-EM map corresponding to the C-terminal His_10_-tag, which may behave as a substrate-like element, was observed near the catalytic site. This observation suggests that opening of the α6-loop may allow a substrate-like polypeptide to approach the catalytic site of BepA, even while the α9-loop remains closed.

### Opening of the BepA α6-loop is required for BepA function on the BAM complex

In the α6-open structure, the opened α6-loop extended from the catalytic site toward the BamA lateral gate. This arrangement raised the possibility that the opened α6-loop interacts with the LptD substrate on BAM and may facilitate its positioning and/or transfer toward BamA. To test this possibility, we examined *in vivo* photo-crosslinking between LptD and BepA using BepA(*p*BPA) variants. LptD-crosslinked products were detected at BepA I168*p*BPA, R190*p*BPA, and M193*p*BPA, which are located on the inner side of the α6-loop and positioned near the BamA lateral gate in the OM, as well as at S63*p*BPA near the catalytic site (**Fig. 4A and B**). These results suggest that the inner side of the opened α6-loop directly interacts with an LptD assembly intermediate. To further examine the BepA–LptD interaction mode, we performed disulfide crosslinking analysis between BepA and LptD. Our previous analysis showed that the edge-strand near the BepA catalytic site is positioned close to Y331 in the N-terminal region of the LptD β-barrel-forming domain (*31*). In contrast, BepA I168C and R190C in the α6-loop did not form clear crosslinking with LptD Y331C but were clearly crosslinked with LptD E391C or V430C (**Fig. 4C**). These cysteine substitutions did not impair BepA or LptD function (**fig. S2D**) (*31*). These results suggest that the α6-loop interacts with a region of LptD distinct from that recognized by the edge-strand.

**Fig. 4.**
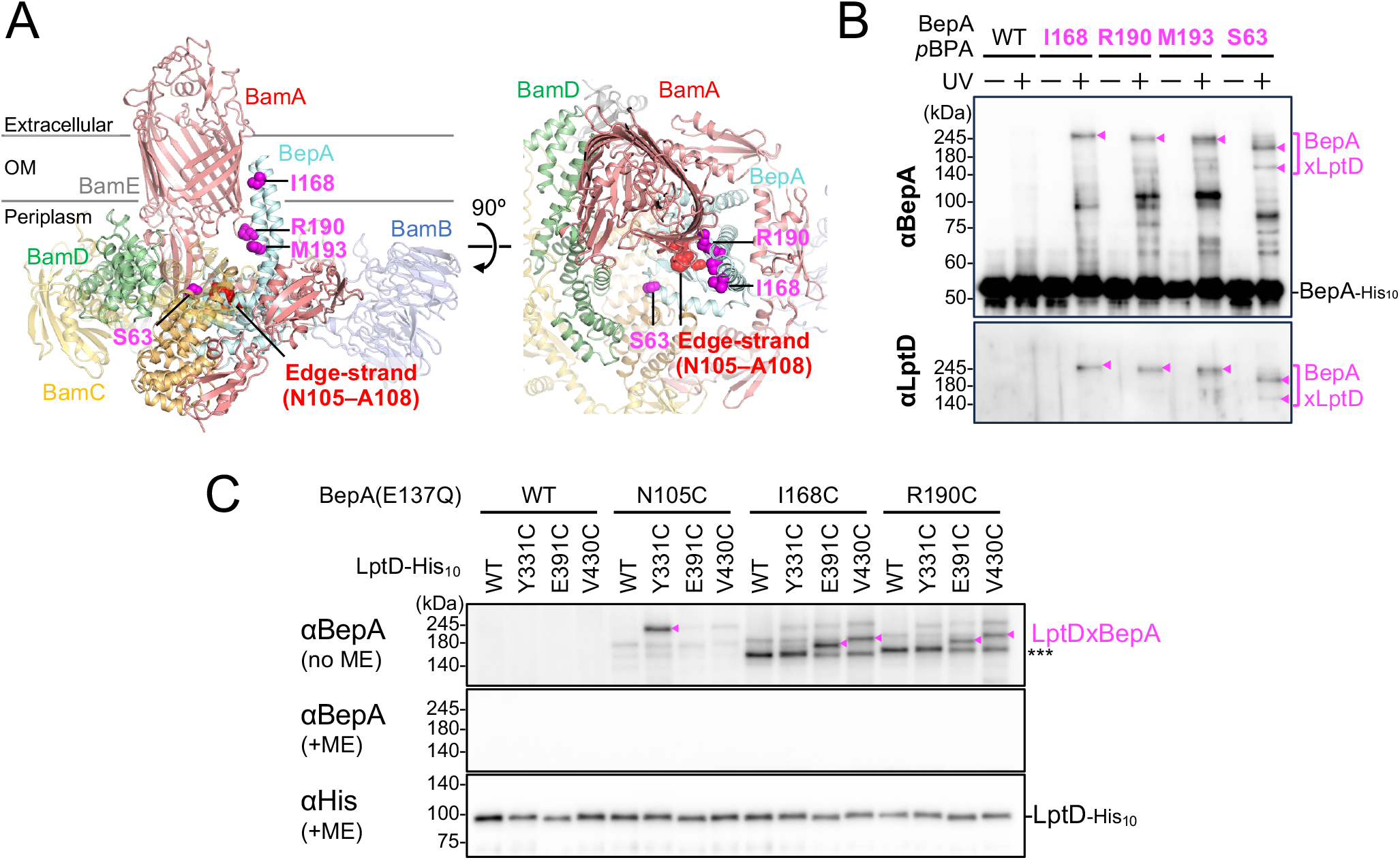
The inner side of the opened BepA α6-loop interacts with an LptD assembly intermediate. (**A**) Mapping of the crosslinking sites on the BAM–BepA α6-open/α9-closed structure (PDB: 27KT). BepA residues crosslinked with LptD are indicated by magenta spheres. The edge-strand is indicated by red spheres. (**B**) *In vivo* photo-crosslinking analysis of BepA. Cells of SN56 (Δ*bepA*) carrying both pEVOL-pBpF and pUC18-*bepA(amb, E137Q)-his_10_* were grown and analyzed using SDS–PAGE followed by immunoblotting using the indicated antibodies as in Fig. 2A. (**C**) Disulfide crosslinking between BepA and LptD. Cells of SN56 (Δ*bepA*) carrying a combination of plasmids encoding WT or a Cys-introduced mutant of BepA and LptD-His_10_ as indicated were grown in LB-medium and induced with 1 mM IPTG for 3 h to express BepA(Cys) and LptD(Cys)-His_10_. Total cellular proteins were acid-precipitated, solubilized with SDS buffer containing NEM (for blocking free thiol groups), and subjected to pull-down with Ni-NTA agarose. The purified proteins were treated with or without 2-mercaptoethanol (ME) and analyzed by 7.5% Laemmli SDS-PAGE and immunoblotting with the indicated antibodies.

To examine the functional role of the α6-loop, we constructed a BepA variant in which the α6-loop was deleted and replaced with a GSGSGS linker (BepAΔα6). As reported previously (*30*, *31*), when LptD is overproduced in Δ*bepA* cells, coexpressed BepA degrades LptD and generates LptD degradation products (**Fig. 5A**). We found that BepAΔα6 apparently had reduced LptD degradation activity compared with wild-type BepA (**Fig. 5A**), suggesting that the α6-loop may contribute to BepA-mediated substrate degradation. We next examined the subcellular localization of the BepAΔα6 variant by fractionation analysis. The catalytically inactive BepA E137Q mutant has been reported to stably interact with BAM and/or substrates, resulting in increased localization to the membrane fraction (*24*, *30*). Consistent with this, wild-type BepA was detected mainly in the soluble fraction, whereas the E137Q mutant was strongly detected in the spheroplast fraction (**Fig. 5B**). In contrast, localization of the BepAΔα6/E137Q mutant to the spheroplast fraction was markedly reduced (**Fig. 5B**). These results suggest that the α6-loop region is important for the localization of BepA near the OM, likely through stable association with BAM and/or its substrates.

**Fig. 5.**
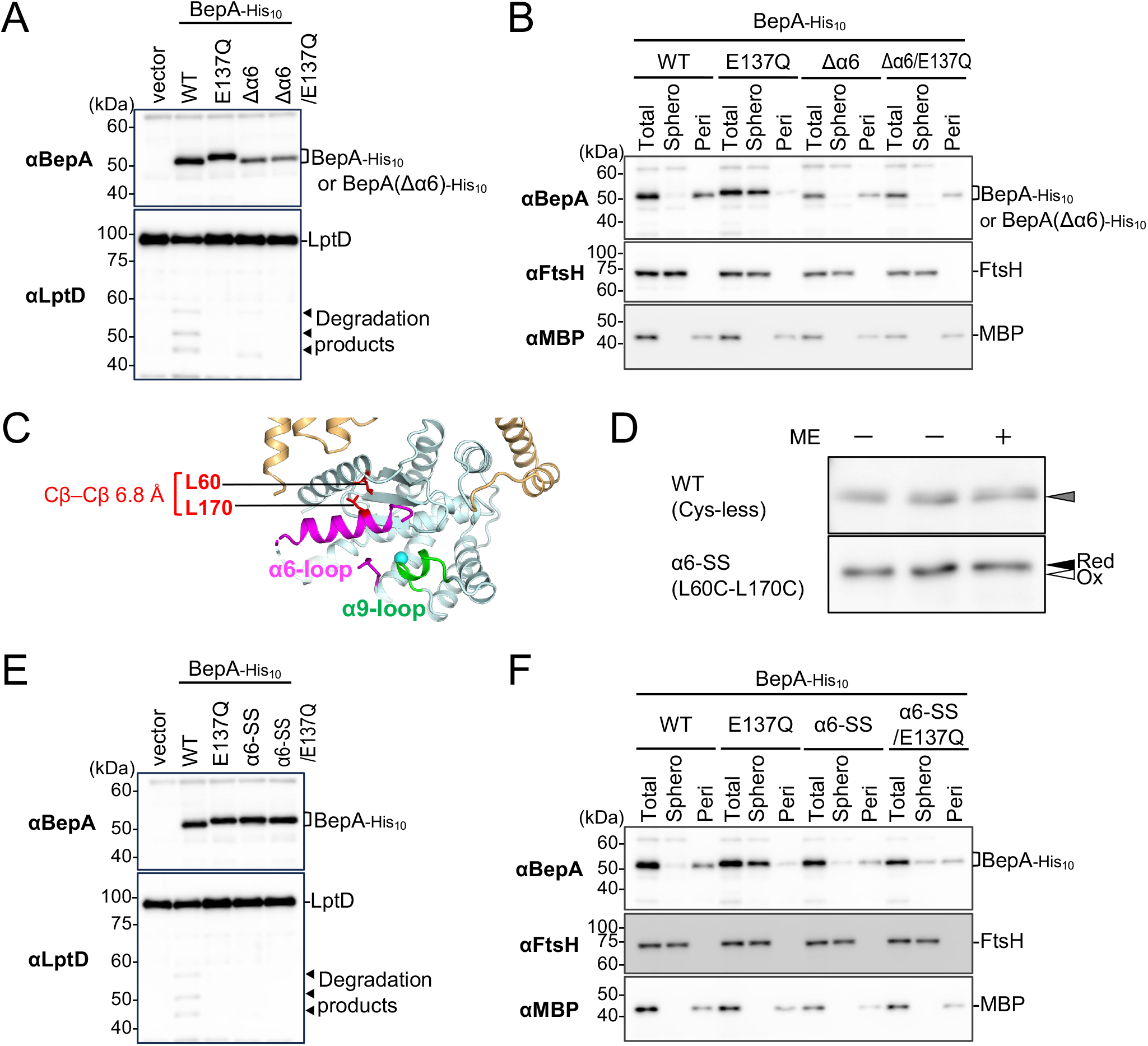
Opening of the BepA α6-loop is required for substrate degradation and membrane-association. (**A**) LptD degradation activity of the BepAΔα6 mutant. Cells of SN56 (Δ*bepA*) carrying pTWV228-*lptD-his_10_* and either pHSG575 or pHSG575-*bepA(mut)-his_10_* plasmids were grown at 30°C in LB-medium until early log phase and induced with 1 mM IPTG for 1 h. Total cellular proteins were acid-precipitated and analyzed by 7.5 or 10% Laemmli SDS-PAGE and immunoblotting with the indicated antibodies. (**B**) Subcellular localization of the BepAΔα6 mutant. Cells of SN56 (Δ*bepA*) carrying pHSG575-*bepA(mut)-his_10_* were grown at 30 °C in LB medium until early log phase, induced with 1 mM IPTG for 2 h to express the indicated BepA variants, and fractionated into spheroplast and periplasmic fractions. Each fraction was analyzed by 7.5 or 10% Laemmli SDS-PAGE and immunoblotting with the indicated antibodies. (**C**) Cysteine substitution sites of the BepA α6-SS mutant. In the crystal structure of BepA (PDB: 6AIT), the α6- and α9-loops are shown in magenta and green, respectively. The introduced cysteine positions are shown as red sticks. (**D**) Disulfide bond formation of α6-SS mutant. Cells of SN56 (Δ*bepA*) carrying pHSG575-*bepA(mut)-his_10_* were grown and induced as in (B). Total cellular proteins were acid-precipitated, treated with or without ME and analyzed by 7.5% Laemmli SDS-PAGE and immunoblotting with anti-BepA antibodies. The oxidized and reduced forms of BepA are indicated as Ox and Red, respectively. (**E**) LptD degradation of the BepA α6-SS mutant. Cells of SN56 (Δ*bepA*) carrying pTWV228-*lptD-his_10_* and either pHSG575 or pHSG575-*bepA(mut)-his_10_* plasmids were grown and analyzed as in (A). (**F**) Subcellular localization of the BepA α6-SS mutant. Cells of SN56 (Δ*bepA*) carrying pHSG575-*bepA(mut)-his_10_* were grown and analyzed as in (B).

However, deletion of the α6-loop region could affect the overall structure of BepA, as the cellular level of BepAΔα6 was slightly lower than that of wild-type BepA. To address this concern, we constructed an α6-SS variant in which the α6-loop was fixed in a closed conformation by introducing cysteine substitutions at L60 and L170, which are located close to each other in the crystal structure (**Fig. 5C**). Formation of an intramolecular disulfide bond in this variant was confirmed in this variant (**Fig. 5D**). We then examined LptD degradation and subcellular localization using the α6-SS variant. Similar to the α6- loop deletion variant, the α6-SS variant showed reduced LptD degradation activity and reduced localization to the spheroplast fraction (**Fig. 5E and F**). Notably, whereas wild-type BepA molecules migrated faster than the E137Q mutant due to C-terminal self-cleavage, the Δα6 and α6-SS variants migrated similarly to their E137Q derivatives, suggesting reduced self-cleavage of the C-terminal His_10_-tag in these mutants (**Fig. 5A and E**). These results suggest that opening of the α6-loop, rather than simply the presence of the α6- loop region, is important for BepA-mediated substrate degradation and BAM-associated function.

### Opening of the BepA α9-loop stabilizes its membrane association

Although opening of the α6-loop is important for BepA function, the α9-loop remained closed in all BAM–BepA structures determined in this study. The NEM accessibility assay also supported the idea that the α9-loop remains relatively closed in living cells. However, for BepA to degrade substrates, the conserved His residue within the α9-loop must be released from coordination with the catalytic Zn ion. We examined how opening of the α9-loop affects the localization of BepA.

We first used the H246A variant that has been reported to exhibit abnormally elevated protease activity and is expected to destabilize the closed α9-loop conformation (*30*). Consistent with this expectation, in the H246A background, BepA S132C and V133C, which are normally poorly modified by thiol-modifying reagents, showed increased NEM modification (**Fig. 6A**). These results suggest that the H246A mutation makes residues covered by the α9-loop more accessible. The extent of NEM modification in the H246A background was comparable to that in the Δα9 variant (**fig. S5**), suggesting that the H246A mutation largely releases the α9-loop from the catalytic site. Fractionation analysis further showed that the H246A variant exhibited increased localization to the spheroplast fraction, even in the absence of the E137Q mutation (**Fig. 6B**).

**Fig. 6.**
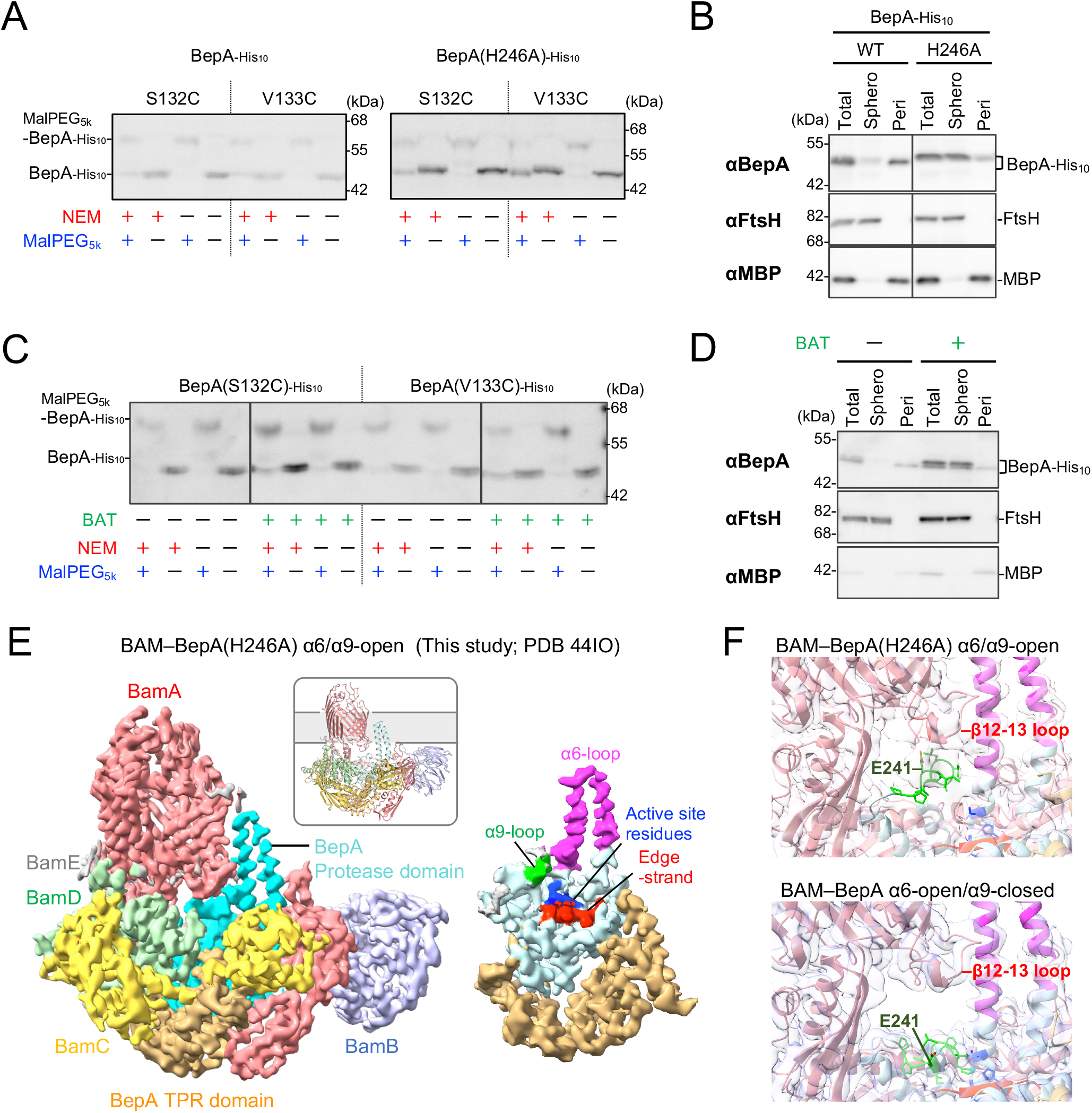
Opening of the BepA α9-loop promotes membrane-associated localization. (**A**) Effect of H246A mutation on the movement of the BepA α9-loop. Cells of SN56 (Δ*bepA*) carrying pHSG575-*bepA(Cys, mut)-his_10_* were grown and analyzed as in Fig. 2E using NEM instead of AMS. (**B**) Effect of H246A mutation on subcellular localization of BepA. Cells of SN56 (Δ*bepA*) carrying pHSG575-*bepA(mut)-his_10_* were grown and analyzed as in Fig. 5B. (**C**) Effect of Batimastat (BAT) treatment on the movement of the BepA α9-loop. Cells of AD2892 (Δ*bepA*, Δ*acrA*) carrying pHSG575-*bepA(Cys)-his_10_* were grown, treated with or without 250 μM BAT, and analyzed as in (A). (**D**) Effect of BAT treatment on subcellular localization of BepA. Cells of AD2892 (Δ*bepA*, Δ*acrA*) carrying pHSG575-*bepA-his_10_* were grown, treated with or without 250 μM BAT, and analyzed as in Fig. 5B. (**E**) Cryo-EM structure of the BAM–BepA α6/α9-open complex. (left) Cryo-EM map and cartoon model of DDM-solubilized BAM– BepA(H246A) complex (PDB: 44IO). BamA, BamB, BamC, BamD, BamE, and the protease and TPR domains of BepA are colored as in Fig. 1C. (right) Cryo-EM map of BepA. The BepA regions are colored as in Fig. 1B. (**F**) Enlarged views of the BepA α9-loop near the BamA β12–β13 loop in both the BAM– BepA(H246A) α6/α9-open structure (PDB: 44IO) and the BAM–BepA α6-open/α9-closed structure (PDB: 27KT). Cryo-EM maps are superimposed onto their structural models.

Because the H246A mutation may cause abnormal behavior of BepA, we next examined whether a similar effect could be observed with wild-type BepA using Batimastat (BAT), a metalloprotease inhibitor that binds to the active-site Zn ion in a substrate-like manner (*36*). In an *acrA* deletion strain, which is expected to increase intracellular BAT accumulation, BAT inhibited BepA-mediated LptD degradation (**fig. S12**), confirming that BAT inhibits BepA protease activity *in vivo*. Under BAT-treated conditions, BepA S132C and V133C showed increased NEM modification, similar to that observed in the H246A variant (**Fig. 6C**). BAT treatment also increased localization of BepA to the spheroplast fraction (**Fig. 6D**). Together, these results suggest that opening of the α9-loop promotes membrane. association of BepA, likely reflecting stabilization of its interaction with BAM and/or stalled substrates. Thus, α6-loop opening and α9-loop opening appear to represent separable steps that together promote BepA function at BAM.

To understand how α9-loop opening stabilizes the BAM–BepA interaction, we introduced the H246A mutation into BepA and determined the structure of the BAM–BepA(H246A) complex in the presence of BAT (**Fig. 6E**; **fig. S13)**. The overall architecture of this BAM–BepA(H246A) complex was similar to that of the α6-open/α9-closed BAM–BepA structure. Although no clear map corresponding to BAT was observed in this structure, the α9-loop was open as expected. In this α9-open conformation, the active site and edge-strand of BepA are exposed and accessible to a substrate polypeptide. In addition, the opened α9-loop directly contacts the periplasmic β12–13 loop region of BamA (**Fig. 6F**). Consistent with this structural observation, BepA E241*p*BPA in the α9-loop was crosslinked to BamA in living cells (**Fig. 2A**), suggesting that the α9-loop can adopt an open conformation and contact BamA *in vivo*. These results suggest that interaction between the opened α9-loop and the BamA β12–β13 loop may stabilize the BAM– BepA complex and may also contribute to regulation of α9-loop opening for BepA activation.

### Discussion

BepA is a periplasmic metalloprotease that contributes to OM integrity, and is thought to mediate the triage of LptD assembly intermediates to promote either their normal assembly or proteolytic removal at the BAM complex (*24*). Previous biochemical and structural studies indicated that the α6- and α9-loops, which cover the proteolytic active site, may undergo dynamic conformational changes to mediate this triage process (*31–33*). However, it has remained unclear how these loop regions undergo conformational changes to regulate BepA function. In this study, we determined multiple distinct BAM–BepA structures, including α6/α9-closed, α6-open/α9-closed and α6/α9-open states (**Figs. 1, 3, and 6**), revealing the interaction mode between BAM and BepA and multiple conformational states of BAM-bound BepA. We demonstrated through *in vivo* photo-crosslinking and accessibility analyses that these BAM–BepA architectures can form *in vivo* and that the α6- and α9-loops undergo dynamic and regulated conformational changes in living cells (**Figs. 2 and 6**). Furthermore, biochemical analyses demonstrated the physiological importance of α6- and α9-loop movements for efficient BepA function (**Figs. 4–6**).

The multiple BAM–BepA structures determined in this study suggest a stepwise conformational transition in which BepA binds to the BAM complex and sequentially opens its α6-loop and then its α9-loop (**Fig. 7**). In the nanodisc-reconstituted BAM–BepA structure, the overall BAM architecture is most similar to that of the BepA-free BAM complex (**Fig. 7A**). In this structure, BepA is located relatively far from the OM and adopts an α6/α9-closed conformation similar to that observed in the previously reported BepA crystal structure. Thus, this structure may represent an early BAM-bound state of BepA (state 1). Then, the BAM–BepA complex may transition to state 2, with the α6/α9-closed BepA moving into the interior of BAM. This intermediate may subsequently shift to state 3, in which BepA adopts an α6-open/α9-closed conformation and is inserted more deeply within BAM. In state 3, the deeply inserted BepA interacts more extensively with BAM than in states 1 and 2, suggesting that BepA may interact more stably with BAM in this state. This stabilized interaction with BAM might facilitate α6-opening of BepA during the transition from state 2 to state 3. In the presence of an aberrantly stalled substrate, BAM–BepA may then adopt an α6/α9-open conformation resembling that of the BAM–BepA(H246A) structure (state 4), in which opening of the α9-loop further stabilizes the BepA–BamA interaction and enables substrate degradation. In state 3, the BamA POTRA domains in the BAM complex adopt a more expanded arrangement. This expansion may allow BAM to adopt a conformation suited for the assembly of large OMP substrates such as LptD. Thus, structural rearrangements of BAM induced by BepA binding may also facilitate LptD maturation.

**Fig. 7.**
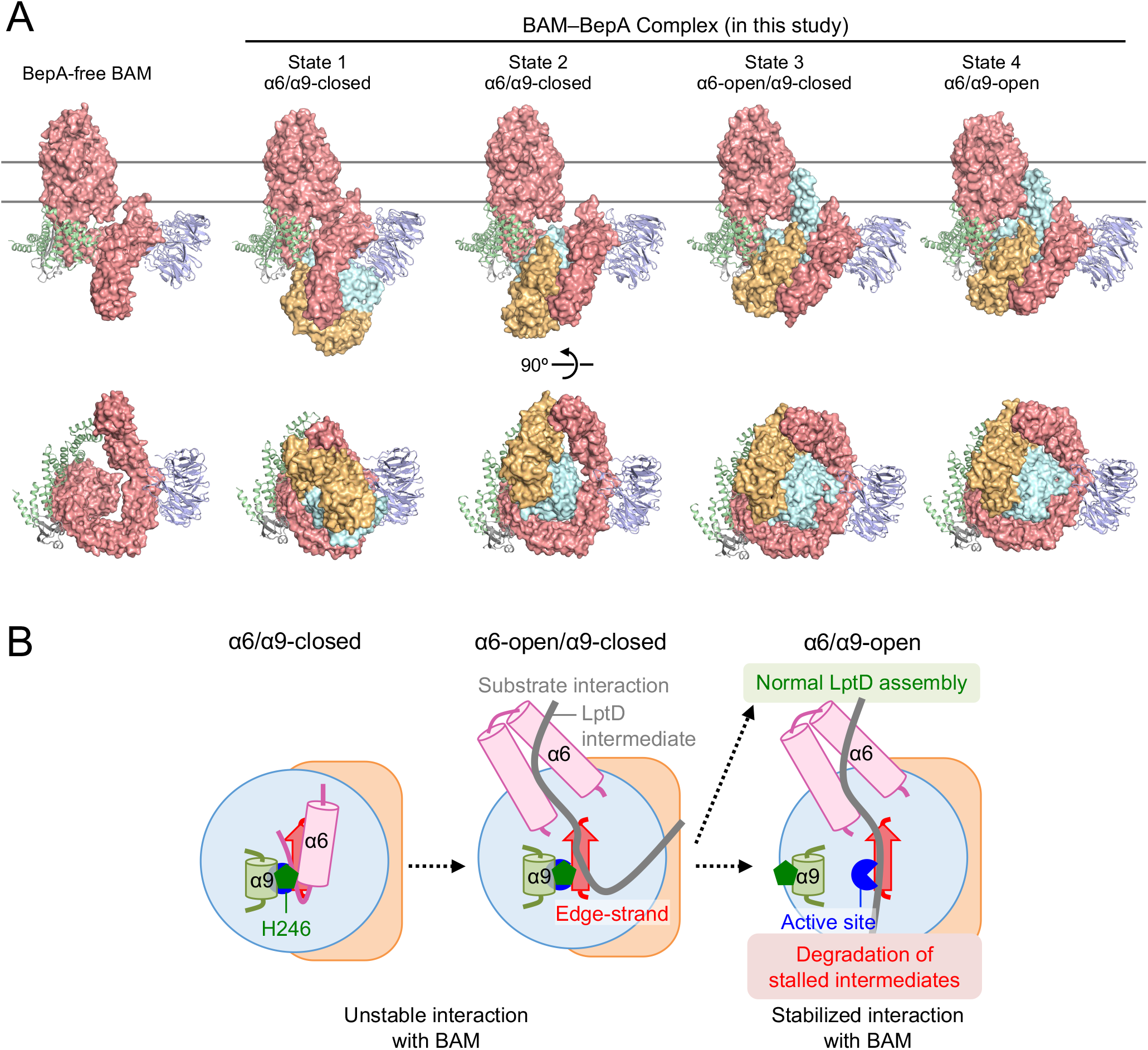
Model for BepA-mediated substrate triage on the BAM complex. (**A**) Cryo-EM structures of the BAM–BepA complex. The BepA-free BAM structure is a previously reported BAM structure (PDB: 9CNW). BAM–BepA structures were determined in this study; α6/α9-closed state 1 (PDB: 27LP), α6/α9-closed state 2 (PDB: 27LF), α6-open/α9-closed state 3 (PDB: 27KT), and α6/α9-open state 4 (PDB: 44IO). BamA, and the BepA protease and TPR domains are shown as red, pale cyan and orange surfaces, respectively. BamB, BamD and BamE are shown as light blue, light green, and gray cartoons, respectively. BamC is omitted for clarity. (**B**) Proposed model of BepA-mediated substrate triage on the BAM complex. Sequential opening of the α6- and α9-loops enables BepA to interact with LptD intermediates and direct them toward either productive maturation or proteolytic degradation. The opened α6-loop supports substrate interaction, whereas α9-loop opening stabilizes BAM association and promotes degradation of aberrantly stalled intermediates.

α6-loop opening may not only rearrange the BAM complex but also allow BepA to directly contact substrate intermediates. In the α6-open BAM–BepA structure, the active site, the edge-strand and the inner surface of opened α6-loop region faced the substrate recognition sites of BAM, the BamA lateral gate and BamD (**Fig. 4**). This arrangement may be suitable for guiding substrates toward the BamA lateral gate. Consistent with this model, residues in these regions were crosslinked to LptD assembly intermediates (**Fig. 4**) (*31*). Thus, the opened α6-loop region may bind to substrate intermediates and may function as a chaperone-like element that physically supports β-barrel formation of substrates near and within the OM. In this state, substrate interaction with the edge-strand may remain limited because the edge-strand is partially covered by the α9-loop (**Fig. 7B**; α6-open/α9-closed). Such transient and weak interactions may enable BepA to act in a chaperone-like manner by repeatedly engaging and releasing LptD assembly intermediates during their maturation.

In contrast, when a substrate becomes aberrantly stalled on the BAM–BepA complex, α9-loop opening may enable the substrate to interact tightly with the edge-strand, leading to its degradation (**Fig. 7B**; α6/α9-open). Initiation of BepA-mediated degradation of stalled LptD assembly intermediates appears to be slower than that of BepA-mediated maturation (*24*). This delay raises the possibility that the generation of aberrantly stalled substrates leads to α9-loop opening and the resulting substrate degradation. Consistent with this idea, a recent study using AlphaFold3 suggested that peptides derived from substrate OMPs can bind to the BepA edge-strand when the α9-loop adopts an open conformation (*37*). In addition, the interaction of the α9-loop with the BamA β12–13 loop may stabilize the α9-open state. Thus, the combination of substrate stalling and the α9-loop–BamA interaction may promote and stabilize α9-opening, enabling BepA to selectively degrade aberrantly stalled substrates on BAM.

Although BepA functions near the BAM complex, most BepA is localized in the periplasmic fraction under physiological conditions (**Fig. 5**). However, it remains unclear when BepA associates with the BAM complex and at which stage and how it dissociates. SurA is another BAM-interacting factor that delivers unfolded OMPs to BAM. SurA mainly contacts BamA, BamB, and BamE and is located at the periphery of the periplasmic side of the BAM complex (*11–13*), whereas BepA mainly contacts BamA, BamC, and BamD and is inserted more deeply into the interior of BAM (**Fig. 3**). These distinct binding sites raise the possibility that SurA and BepA can transiently coexist on BAM, potentially allowing substrate transfer between them. Recently, it was reported that BepA became enriched at the OM in a Δ*surA* background, possibly reflecting the accumulation of misfolded or stalled OMPs (*38*). Taken together, these observations suggest that BepA may associate with BAM either during substrate handover from SurA or after substrate delivery to BAM. In a recently reported cryo-EM structure of a BAM-bound LptD/E assembly intermediate, LptE is inserted into the forming β-barrel of LptD (*19*), indicating that this structure represents a later step of LptD maturation. Notably, the BamA POTRA domains in this structure adopt a more compact arrangement than those in the α6-open BAM–BepA structure (**fig. S14)**, raising the possibility that BepA may dissociate from BAM before this maturation step. Because persistent BepA binding within BAM might interfere physically and/or functionally with the assembly of large OMP substrates, BepA dissociation may facilitate subsequent LptD maturation. Consistent with this model, LptM acts at a later step of LptD maturation than BepA (*28*).

Whether BepA acts on substrates other than LptD remains an important question. The suppression of the Δ*bepA* phenotype by LptE-overexpression strongly suggests that LptD is a major physiological substrate of BepA (*24*). Nevertheless, BepA has also been implicated in BamA degradation in a Δ*surA* background (*24*, *34*), and a recent study reported that BepA can degrade OmpX-derived peptides *in vitro* (*37*). These observations raise the possibility that BepA contributes more broadly to OMP quality control. It also remains unclear whether BepA acts on soluble periplasmic substrates.

OMP biogenesis and LPS transport are essential processes in Gram-negative bacteria and have therefore attracted increasing attention as antibacterial targets. Several small-molecule inhibitors have already been developed, and phage- or bacterium-derived peptides that inhibit these pathways have also been identified, highlighting the therapeutic potential of targeting OM biogenesis (*39–42*). Because BepA is nonessential but maintains OM integrity, its inhibition may increase OM permeability and enhance antibiotic susceptibility with reduced selective pressure for resistance to such inhibitors. Although BAT appears to bind to and inhibit BepA, the binding affinity is rather low. Our BAM–BepA structures provide structural insights for the future development of new BepA-targeting inhibitors. Taken together, our BAM– BepA structures advance the mechanistic understanding of periplasmic protease-mediated OMP quality control and may guide future efforts to develop antibacterial agents that disrupt proper OMP assembly in Gram-negative bacteria.

## Methods

### Bacterial strains and plasmids

The *E. coli* strains and plasmids used in this study are listed in tables S1 and S2, respectively. Details of the mutant strains and plasmids constructed in this study are described in the Construction of mutant strains and Plasmid construction sections. Strains and plasmids generated in this study are available from the corresponding authors upon request.

### Media and bacterial cultures

The cells were grown in LB medium (Nacalai Tesque), with 50 or 100 µg/mL ampicillin (Amp), 20 µg/mL chloramphenicol (Cm), 25 µg/mL kanamycin (Km), 25 µg/mL tetracycline (Tet), and/or 50 µg/mL spectinomycin (Spc) added as appropriate for growing the plasmid-bearing cells and selecting transformants and transductants. Bacterial growth was monitored using a Mini photo 518R (660 nm; TAITEC Co.).

### Construction of mutant strains

AD2892 (AD16, Δ*bepA*, Δ*acrA*::*kan*) was constructed by transducing Δ*acrA*::*kan* from JW0452 (*43*) to SN56 (*24*). RM4676 (HM1742, Δ*bepA*, *purC80*::Tn*10*) was constructed by transducing Δ*bepA*, *purC80*::Tn*10* from RM2098 (*31*) to HM1742 (*44*). RM4748 (HM1742, *purC80*::Tn*10*, *kan araC-P_araBAD_*-*bamA*) was constructed by transducing *kan araC-P_araBAD_-bamA* from RM4714 (*13*) to RM4676.

### Plasmid construction

BAM–BepA expression plasmids used in this study were constructed as described below. In all of these plasmids, the encoded Bam components and BepA retained their native signal sequences. pRM1596 (pJH113-*bamA(P47C, C690S, C700S)/bamB/bamC/bamD/bamE-his_8_*) and pRM1598 (pJH113-*bamA(C690S, C700S, P779C)/bamB/bamC/bamD/bamE-his_8_*) were constructed from pRM1261 (*13*) by site-directed mutagenesis. pRM1752 (pJH113-*bamA(C690S, C700S, P779C)/bamB/bamC/bamD/bamE/his_10_-bepA(E137Q, A180C)-his_10_*) and pRM1763 (pJH113-*bamA(P47C, C690S, C700S)/bamB/bamC/bamD/bamE/his_10_-bepA(E137Q, F404C)-his_10_*) were constructed by *in vitro* recombination using the In-Fusion HD Cloning Kit. For pRM1752, a vector fragment containing *bamA(C690S, C700S, P779C)/bamB/bamC/bamD/bamE* was amplified from RM1598 using the pBAM-F2 (5’-GATCCTCTAGAGTCGACCTG-3’) and pBAM-R2 (5’-TTAGTTACCACTCAGCGCAGG-3’) primers, and a *his_10_-GGSG-bepA(E137Q, A180C)-his_10_* fragment (N-terminal His_10_-GGSG tag was inserted after the signal sequence of BepA) was amplified from pRM1615 using the BAM-bepA-F (5’-CTGAGTGGTAACTAACCAGAAATACAGGATAGAGG-3’) and BAM-bepA-R (5’-CGACTCTAGAGGATCGGATCCCCGGGTACTTAATG-3’) primers. For pRM1763, a vector fragment containing *bamA(P47C, C690S, C700S)/bamB/bamC/bamD/bamE* was amplified from RM1596 using the pBAM-F2 and pBAM-R2 primers, and a *his_10_-GGSG-bepA(E137Q, F404C)-his_10_* fragment was amplified from pRM1617 using the BAM-bepA-F and BAM-bepA-R primers. The corresponding *bamA–bamE* and *bepA* fragments were ligated by *in vitro* recombination. pRM1949 (pJH113-*bamA(P47C, C690S, C700S)/bamB/bamC/bamD/bamE/his_10_-bepA(E137Q, F404C)*) was constructed from pRM1763 by site-directed mutagenesis. pRM2111 (pJH113-*bamA(P47C, C690S, C700S)/bamB/bamC/bamD/bamE/his_10_-bepA(E137Q, H246A, F404C)*) was constructed from pRM1949 by site-directed mutagenesis.

pUC18-*bepA(E137Q, amb)-his_10_* and pUC18-*bepA(amb)-his_10_* plasmids were constructed from pUC-*bepA(E137Q)-his_10_* (*24*) and pUC-*bepA-his_10_* (*24*), respectively, by site-directed mutagenesis. pYD552 (pHSG575-*bepA-his_10_*) and pYD556 (pHSG575-*bepA(E137Q)-his_10_*) were constructed by subcloning the *EcoRI–KpnI* fragments carrying *bepA-his_10_* or *bepA(E137Q)-his_10_* from pUC-*bepA-his_10_* or pUC-*bepA(E137Q)-his_10_*, respectively, into the same sites of pHSG575 (NBRP). pHSG575-*bepA(Cys)-his_10_* plasmids were constructed from pYD552 by site-directed mutagenesis. pRM1474 (pHSG575-*bepA(*Δ*α6)-his_10_*) and pRM1476 (pHSG575-*bepA(*Δ*α6, E137Q)-his_10_*) were constructed by subcloning the *EcoRI– HindIII* fragments carrying *bepA(*Δ*α6)-his_10_* or *bepA(*Δ*α6, E137Q)-his_10_* from pRM1365 or pRM1385, respectively, into the same sites of pHSG575. pSTD1636 (pHSG575-*bepA(L60C, L170C)-his_10_*) and pSTD1640 (pHSG575-*bepA(L60C, L170C, E137Q)-his_10_*) plasmids were constructed from pYD552 and pYD556, respectively, by sequential site-directed mutagenesis. pSTD1519 (pHSG575-*bepA(H246A)-his_10_*) was constructed from pYD552 by site-directed mutagenesis. pSTD1525 (pHSG575-*bepA(S132C, H246A)-his_10_*) and pSTD1527 (pHSG575-*bepA(V133C, H246A)-his_10_*) plasmids were constructed from pSTD1519 by site-directed mutagenesis. For construction of pSTD1570 (pHSG575-*bepA(*Δ*α9)-his_10_*), the Δ*α9* mutation (*33*) was introduced into pTH-bepA-his10 (*24*) by site-directed mutagensis, and then the EcoRI-SalI fragment of the resulting plasmid, which contained pHSG575-*bepA(*Δ*α9)-his_10_*, was recloned into the same site of pHSG575. pSTD1625 (pHSG575-*bepA(S132C,* Δ*α9)-his_10_*) and pSTD1627 (pHSG575-*bepA(V133C,* Δ*α9)-his_10_*) were constructed from pSTD1570 by site-directed mutagenesis. pSTD689-*bepA(Cys)* and pSTD689-*bepA(E137Q, Cys)* plasmids were constructed from pRM290(*30*) and pRM29(*30*), respectively, by site-directed mutagenesis. pRM866 (pSTD689-*bepA-his_10_*) and pRM868 (pSTD689-*bepA(E137Q)-his_10_*) were constructed by subcloning the *EcoRI–SalI* fragments carrying *bepA-his_10_* or *bepA(E137Q)-his_10_* from pUC-*bepA-his_10_* or pUC-*bepA(E137Q)-his_10_*, respectively, into the same sites of pSTD689(*45*). pSTD689-*bepA(E137Q, Cys)-his_10_* plasmids were constructed from pRM868 by site-directed mutagenesis. pRM1612 (pSTD689-*his_10_*-*bepA(E137Q)-his_10_*) was constructed by introducing the *his_10_-ggsg*-tag sequence into pRM868 by In-Fusion cloning using an inverse PCR-amplified vector fragment generated with primers H-bepA-F (5’-CATCACCATCACGGCGGTTCTGGCGCAGACACCTTGCCGGATATG-3’) and H-bepA-R (5’-GCCGTGATGGTGATGATGGTGGTGATGGTGATGGCTGTCGGCAAACGCCGG-3’). pSTD689-*his_10_*-*bepA(E137Q, Cys)-his_10_* plasmids were constructed from pRM1612 by site-directed mutagenesis. pUC118-*bamA(Cys)* plasmids were constructed from pRM864(*13*) by site-directed mutagenesis.

### Immunoblotting analysis

Acid-denatured proteins, prepared as described in the figure legends for each experiment, or purified proteins were solubilized in SDS-sample buffer (62.5 mM Tris–HCl [pH 6.8], 2% SDS, 10% glycerol, and 0.01% [w/v] bromophenol blue) with or without 10% β-mercaptoethanol (ME). The samples were incubated at 98 °C for 5 min, separated by SDS‒PAGE, and electroblotted onto PVDF membranes (Merck Millipore). The membranes were first blocked with 1% skim milk in phosphate-buffered saline with Tween 20 (PBST) and incubated with anti-BepA (*24*) (1/5,000 or 1/10,000 dilution), anti-LptD (*24*) (1/50,000), anti-BamA (*46*) (1/100,000), anti-BamB (*46*) (1/10,000), anti-BamD (*46*) (1/10,000), anti-FtsH (*47*) (1/10,000), or anti-MBP (*48*) (1/200,000) antibodies. After washing with PBST, the membrane was incubated with a horseradish peroxidase (HRP)-conjugated secondary antibody (goat anti-rabbit IgG (H + L)-HRP conjugate; Bio-Rad, catalog no. 1706515; 1/5,000) in PBST. After washing with PBST, the proteins were visualized using detection reagents (ECL^TM^ Western Blotting Detection Reagents (Cytiva), ECL^TM^ Prime Western Blotting Detection Reagents (Cytiva), Chemi-Lumi One (Nacalai Tesque) or Chemi-Lumi One Super (Nacalai Tesque)) and chemiluminescence image analyzers (LAS4000 mini lumino-image analyzer (Cytiva) and FUSION Solo S (VILBER)).

### Purification of the BAM–BepA complexes

BL21(DE3) (Novagen) cells harboring pRM1949 (**figs. S1A and S6A**), pRM1763 (**figs. S3A and S9A**), pRM1752 (**fig. S11A**), or pRM2111 (**fig. S13A**) were inoculated into LB medium supplemented with 0.4% glucose and 50 μg/mL ampicillin and incubated at 37 °C for 14–16 h. The overnight culture was then inoculated with LB medium containing 50 μg/mL ampicillin and grown at 30 °C until the OD_600_ reached 0.8–0.9. Isopropyl β-D-thiogalactopyranoside (IPTG) was added at a concentration of 0.5 mM. The cells were further cultured at 37 °C for 1.5 h to induce the BAM–BepA complex expression. Cells were collected by centrifugation, suspended in buffer (20 mM Tris–HCl [pH 8.0], 150 mM NaCl, 1 mM EDTA-Na [pH 8.0], and 0.1 mM PMSF), and disrupted using a Microfluidizer Processor M-110EH at 100 MPa (Microfluidics International). After removing the unbroken cells and protein aggregates by centrifugation, the supernatant was subsequently ultracentrifuged at 186,000 x g for 60 min at 4 °C (Beckman 45Ti rotor) to isolate the membrane fraction. The membrane fraction was resuspended in solubilization buffer (50 mM Tris–HCl [pH 8.0], 150 mM NaCl, 1% [w/v] n-dodecyl β-maltoside [DDM], 20 mM imidazole-HCl [pH 8.0], and 0.1 mM PMSF) and solubilized by gentle stirring at 4 °C for 60 min. After removal of the insoluble fraction by ultracentrifugation at 186,000 x g for 30 min at 4 °C (Beckman 45Ti rotor), the supernatant was mixed with 2.5 ml of Ni-NTA agarose resin (QIAGEN), pre-equilibrated with solubilization buffer, and gently stirred at 4 °C for 60 min. Next, 2.5 mL of wash buffer (50 mM Tris–HCl [pH 8.0], 150 mM NaCl, 0.05% [w/v] DDM, 50 mM imidazole-HCl [pH 8.0], and 0.1 mM PMSF) was added to the column three times. Then, 2.5 mL of elution buffer (50 mM Tris–HCl [pH 8.0], 150 mM NaCl, 0.05% [w/v] DDM, 500 mM imidazole-HCl [pH 8.0], and 0.1 mM PMSF) was added to the column six times. The eluted fractions containing BamA-crosslinked to His_10_-BepA, BamB, BamC, BamD, and BamE were collected and concentrated using an Amicon Ultra 50K NMWL (Merck Millipore). In the case of the BAM– BepA(H246A) complex, the purified proteins were treated with 300 μM BAT on ice for 30 min. After ultracentrifugation at 106,400 x g for 30 min at 4 °C (Himac S55A2 rotor), the concentrated sample was loaded onto a Superdex^TM^ 200 Increase 10/300 GL column (Cytiva) equilibrated with SEC buffer (50 mM Tris–HCl [pH 8.0], 150 mM NaCl, 0.02% [w/v] DDM, and 0.1 mM PMSF). The fractions eluted with the target proteins were collected and further concentrated using an Amicon Ultra 50K NMWL.

### Nanodisc reconstitution of the BAM–BepA complex

The purified BAM–BepA complex was mixed with MSP1E3D1 (*49*) and E. coli phospholipids (Avanti) at a molar ratio of 1:4:400 in SEC buffer. The mixture was incubated at 4°C for 1 hour. After addition of Bio-Beads SM2 (Bio-Rad), the mixture was gently rotated overnight at 4°C to initiate detergent removal and nanodisc formation. Next, the mixture was filtered using Centrifugal Filters PVDF 0.22 μm (Millipore). The filtrate was then ultracentrifuged at 186,000 x g for 30 min at (himac S55A2 rotor). The resulting supernatant was applied to a Superose 600 10/300 GL column (Cytiva) or a Superdex 200 10/300 GL column preequilibrated with SEC buffer without DDM. Fractions were analyzed by SDS–polyacrylamide gel electrophoresis (SDS-PAGE) and native PAGE to confirm nanodisc reconstitution. Fractions containing the BAM–BepA-reconstituted nanodiscs were pooled and concentrated using an Amicon Ultra 50 K NMWL (Merck Millipore).

### Cryo-EM grid preparation

Quantifoil holey carbon grids (Cu R1.2/R1.3, 300 mesh) were glow-discharged at 7 Pa and 10 mA for 10 seconds using a JEC-3000FC sputter coater (JEOL) before sample application. For the nanodisc-reconstituted BAM–BepA-His_10_ complex, Quantifoil holey carbon grids (Cu R1.2/R1.3, 300 mesh) were glow-discharged for 45 s at 20 mA using a GloQube (Quorum) instrument. A 3-μL aliquot of the DDM-solubilized BAM–BepA complexes (∼15 mg/mL) or the nanodisc-reconstituted BAM–BepA complexes (3∼4 mg/mL) was applied onto the grids, which were blotted for 2, 3, or 4 seconds at 100% humidity and 8 °C with a blot force of 10 or 4, and then plunged into liquid ethane using a Vitrobot Mark IV (Thermo Fisher Scientific).

### Cryo-EM data collection and processing

For the nanodisc-reconstituted BAM–BepA complex, the BAM–BepA-His complex, the β-barrel–α6-closslinked BAM–BepA-His complex, and the BAM–BepA(H246A) complex, cryo-EM datasets were collected using a CRYO ARM 300 transmission electron microscope (JEOL) operated at an accelerating voltage of 300 kV, equipped with a cold-field emission gun, an in-column Omega-type energy filter, and a Gatan K3 camera (Gatan) at SPring-8. Images were collected at a nominal magnification of 60,000×, corresponding to a calibrated pixel size of 0.752 Å/pixel, 50 frames per image with a total exposure dose of 50 e−/Å2 using a SerialEM (version 4.2.0)(*50*).

For the BAM–BepA complex, cryo-EM datasets were collected using a Glacios Cryo-TEM instrument operated at an accelerating voltage of 200 kV and equipped with a Falcon4i direct electron detector (Thermo Fisher Scientific) at SPring-8. Images were collected at a nominal magnification of 190,000×, corresponding to a calibrated pixel size of 0.74 Å/pixel, 50 frames per image with a total exposure dose of 55 e−/Å2, via the EPU software (version 3.13.0.10277).

For the nanodisc-reconstituted BAM-BepA-His complex, the cryo-EM dataset was collected on a Krios G4 Cryo-TEM instrument operated at an accelerating voltage of 300 kV and equipped using a Selectris X energy filter and Falcon4i direct electron detector (Thermo Fisher Scientific) at National Research and Innovation (BRIN) in Indonesia. Images were collected at a nominal magnification of 165,000×, corresponding to a calibrated pixel size of 0.76 Å/pixel, 40 frames per image with a total exposure dose of 30 e−/Å2, using the EPU software (version 3.8.1.7603; Thermo Fisher Scientific).

Data processing was performed using the CryoSPARC v4.5.3–v5.0.6 software platform(*51*). Cryo-EM data-processing for various BAM–BepA complexes was performed by the procedures shown in **figs. S1D, S3D, S6D, S9D, S11D and S13D.** The acquired movies were aligned using patch motion correction, and the contrast transfer function (CTF) parameters were estimated using the Patch CTF estimation. The particles were initially auto-picked from 200, 500 or 1000 micrographs using the Blob Picker, and the resulting particles were subjected to 2D classification. The obtained 2D class averages were then used as templates for automatic particle picking with the Template Picker. Particles were extracted with a downsampling via Fourier cropping and subjected to 2D classification. Next, these selected particles were subjected to ab initio 3D reconstruction to generate initial 3D models. The particles were then subjected to several rounds of heterogeneous refinement to remove junk particles using the initial models as references. The best 3D class particles were re-extracted at the original pixel size. These particles were subjected to global/local CTF refinement, reference-based motion correction and Non-Uniform (NU) refinement, which resulted in final cryo-EM maps. For the dataset of the BAM–BepA complex, to analyze structural heterogeneity, the refined 183,274 particles were subjected to two rounds of 3D classification, one without alignment and another with alingment, in CryoSPARC. Each particle subset was subjected to NU refinement, resulting in two final maps: an α6/α9-closed BAM–BepA complex and an α6-open/α9-closed BAM–BepA complex (**fig. S3D**).

### Model building and refinement

The BAM complex model from the SurA–BAM structures (PDB ID: 25FQ) and the BepA model from the AlphaFold2/3-predicted model (*52*, *53*) were docked into the obtained cryo-EM maps using UCSF Chimera X(*54*). The fitted structures were manually adjusted in Coot (version 0.9.8.96) (*55*), followed by further refinements using the phenix.real_space_refine in PHENIX (version1.21.2) (*56*). The data processing and refinement statistics are summarized in table S3. The molecular model and cryo-EM map were visualized using PyMOL (version 2.6.2; https://pymol.org/) or UCSF ChimeraX (version 1.9).

### *In vivo* photo-crosslinking analysis

pEVOL-pBpF expresses an evolved tRNA/aminoacyl-tRNA synthetase pair that enables the *in vivo* incorporation of *p*BPA into an amber codon site of a target protein via *amber* suppression (*57*). UV irradiation to a cell expressing a *p*BPA-incorporated protein induces the formation of covalent crosslinking products between *p*BPA in the target protein and a nearby protein, which allows for the detection of their *in vivo* interaction (*58*, *59*). Cells were grown at 30 °C in LB medium supplemented with 0.5 mM *p*BPA and 0.02% arabinose until the early log phase, followed by induction with 1 mM IPTG for 1 h. Half of the cell cultures was transferred to a Petri dish and UV-irradiated at 4 °C for 10 min using a B-100AP UV lamp (365 nm; UVP, LLC.) at a distance of 4 cm. The other half was placed on ice as a non-UV-irradiated samples. Total cellular proteins were precipitated with 5% trichloroacetic acid (TCA), washed with acetone, solubilized in SDS buffer (50 mM Tris-HCl [pH 8.0], 1% SDS, and 1 mM EDTA). After 33-fold diluted with Triton buffer (50 mM Tris-HCl [pH 8.0], 150 mM NaCl, 2% Triton X-100, and 0.1 mM EDTA) and clarification, samples were subjected to pull-down with Ni-NTA Agarose (QIAGEN). The isolated proteins were solubilized in SDS-sample buffer containing ME, boiled at 98°C for 5 min, and analyzed by SDS-PAGE and immunoblotting analysis.

### AMS/NEM accessibility assay

Cells of SN56 (Δ*bepA*) carrying pHSG575-*bepA(Cys)-his_10_* were grown at 30 °C in LB-medium and induced with 1 mM IPTG for 2 h to express BepA derivatives. Cells were harvested, resuspended in 100 µL of spheroplast buffer (20% sucrose, 30 mM Tris-HCl [pH 8.1]), and converted to spheroplasts by adding 10 µL of 1 mg/mL lysozyme in 0.1 M EDTA (pH 7.0), followed by incubation at 0 °C for 30 min. After the addition of 2.2 µL of 1 M MgCl_2_, 2.2 µL of 10 mM TCEP and 1.1 µL of 0.1 M zinc aceteate, 45 µl of the spheropasts were mixed with 5 µL of 10 mM AMS or NEM and incubated at 4°C for 30 min. The AMS/NEM modification was terminated by addition of 5 µL of ME. Total proteins were acid-precipitated, washed with acetone and solubilized in 20 µl of 10 mM Tris-HCl (pH 8.1)-1% SDS-0.2 mM TCEP. Subsequently, 9 µL of the sample was incubated with 1 µl of either 0 or 50 mM MalPEG_5k_ (in DMSO) at 37 °C for 30 min with vigorous mixing. After quencing MalPEG_5k_ by addition of 1 µL of 20 mM NEM followed by incubation at 37 °C for 5 min, samples were mixed with 10 µL of 2xSDS smaple buffer containing 5 mM MalPEG_5k_ (for DMSO-treated samples) or 10% DMSO (for MalPEG_5k_-treated samples). Proteins were analyzed by SDS-PAGE and immunoblotting with the indicated antibodies.

### Disulfide crosslinking

Cells of SN56 (Δ*bepA*) carrying a combination of plasmids encoding WT or a Cys-introduced mutant of BepA and LptD-His_10_ as indicated were grown at 30 °C in LB-medium and induced with 1 mM IPTG for 3 h to express BepA(Cys) and LptD(Cys)-His_10_. Total cellular proteins were acid-precipitated, solubilized with SDS buffer containing 12.5 mM NEM (for blocking free thiol groups). After 33-fold diluted with Triton buffer and clarification, samples and subjected to pull-down with Ni-NTA agarose. The purified proteins were treated with or without ME and analyzed by 7.5% Laemmli SDS-PAGE and immunoblotting with the indicated antibodies.

### Subcellular fractionation

Cells of SN56 (Δ*bepA*) carrying pHSG575-*bepA(wt or mut)-his_10_* were grown at 30 °C in LB-medium and induced with 1 mM IPTG for 2 h to express BepA drivatives. Cells were harvested and resuspended in 400 µL of spheroplast buffer. Cells were converted to spheroplats as described above. After a 90-µL aliquot was removed as the total fraction, the remaining sample was centrifuged, and the supernatant was collected as the periplasmic fraction. The pellet was resuspended in 350 µL of spheroplast buffer supplemented with 1/10 volume of 1 mg/mL lysozyme in 0.1 M EDTA (pH 7.0), followed by addition of 7 µL of 1 M MgCl₂, and used as the spheroplast fraction. Proteins in 90 µL of each fraction were acid-precipitated, washed with acetone, solubilized in SDS sample buffer, and analyzed by SDS-PAGE and immunoblotting with the indicated antibodies.

## Data Availability

The coordinates have been deposited in the Protein Data Bank (PDB), and the corresponding cryo-EM maps have been deposited in the Electron Microscopy Data Bank (EMDB) under the following accession codes: Nanodisc-reconstituted BAM–BepA α6/α9-closed complex, PDB 27LP and EMD-81243; Nanodisc-reconstituted BAM–BepA-His α6/α9-closed, PDB 27ML and EMD-81262; BAM–BepA α6/α9-closed complex, PDB 27LF and EMD-81235; BAM–BepA α6-open/α9-closed complex, PDB 27KT and EMD-81228; BAM–BepA-His α6-open/α9-closed complex, PDB 27MJ and EMD-81260; β-barrel–α6-closslinked BAM–BepA-His 6-open/α9-closed complex, PDB 27MK and EMD-81261; and BAM– BepA(H246A) α6/α9-open complex, PDB 44IO and EMD-82637; respectively.

## Code Availability

This paper does not report original code.

## Acknowledgements

We thank Kayo Abe and Satomi Koshiba for secretarial assistance, Kunihito Yoshikaie for technical support, the scientists of SPring-8 Structural Biology Beamlines for helping with data collection, and National BioResource Project (NBRP)–*E. coli* at National Institute of Genetics, Japan, for providing *E. coli* strains. ChatGPT (OpenAI) was used to improve the English of this manuscript. The cryo-EM experiments were performed at SPring-8 in Japan with the approval of the Japan Synchrotron Radiation Research Institute (proposal nos. 2024B2742, 2025A2725, and 2026A2715), and at National Research and Innovation Agency (BRIN) in Indonesia.

## Funding

This work was supported by JSPS/MEXT KAKENHI (Grant Nos. JP22K15061, JP24KK0138, JP25K00267 to R.M., Grant Nos. JP23K14146, JP25K18422 to H.K., Grant Nos. JP22H02571, JP23K23835 to Y.A., and Grant Nos. JP25K02226, JP22H02567, JP22H02586 to T.T.), JST NEXUS (Nos. JPMJNX25E5), private research foundations (the Institute for Fermentation (Y-2024-02-006 to R.M., Y-2025-2-046 to H.K., and G-2024-2-034 to T.T., Takeda Science Foundation to T.T.), JST SPRING (Grant No. JPMJSP2140 to Y.S.T.), and RIIM-KI JST (Grant No. 76/II.7/HK/2025). This research was partially supported by Platform Project for Supporting Drug Discovery and Life Science Research (Basis for Supporting Innovative Drug Discovery and Life Science Research (BINDS)) from AMED under grant number JP25ama121001.

## Author Contributions

Conceptualization: R.M., and Y. A.

Methodology: R.M., H.K., Y.A., and T.T.

Investigation: R.M., M.K., H.K., T.N., K.S.,Y.S.T., S.A., Y.D., T.S., F.A.L., Y.N., N.D., S.N., H.S., Y.A., and T.T.

Visualization: R.M. and T.T. Supervision: R.M., Y.A. and T.T.

Writing—original draft: R.M. and Y.A.

Writing—review and editing: R.M., Y.A. and T.T.

## Competing interests

The authors declare no competing interests.

## Supplementary materials

figs. S1 to S14 tables S1 to S3

**fig. S1.** Cryo-EM analysis of the nanodisc-reconstituted BAM–BepA complex.

**fig. S2.** Activities of BamA and BepA cysteine derivatives used in this study.

**fig. S3.** Cryo-EM analysis of the nanodisc-reconstituted BAM–BepA-His complex.

**fig. S4.** *In vivo* photo-crosslinking analysis using BepA(*p*BPA) derivatives.

**fig. S5.** Effect of α9-loop deletion on the AMS accessibility of the BepA S132C and V133C derivatives.

**fig. S6.** Cryo-EM analysis of the BAM–BepA complex.

**fig. S7.** Interface between BamA and the BepA protease domain in the α6-open/α9-closed BAM–BepA structure.

**fig. S8.** Mapping of the BAM-crosslinking sites in the BepA TPR domain onto the BAM–BepA α6-open/α9-closed structure.

**fig. S9.** Cryo-EM analysis of the BAM–BepA-His complex.

**fig. S10.** Disulfide crosslinking between BepA and BamA.

**fig. S11.** Cryo-EM analysis of the β-barrel-α6-closslinked BAM–BepA-His complex.

**fig. S12.** Batimastat inhibits the LptD degradation activity of BepA.

**fig. S13.** Cryo-EM analysis of the BAM–BepA(H246A) complex in the presence of Batimastat.

**fig. S14.** Comparison of the BAM–BepA α6-open/α9-closed and BAM–LptDE structures.

**table S1.** Strains used in this study.

**table S2.** Plasmids used in this study.

**table S3.** Cryo-EM data collection and refinement statistics.

## Notes

### Competing Interest Statement

The authors have declared no competing interest.

## Reference

1. H. Nikaido, Molecular Basis of Bacterial Outer Membrane Permeability Revisited. Microbiol. Mol. Biol. Rev. 67, 593–656 (2003).

2. A. Konovalova, D. E. Kahne, T. J. Silhavy, Outer Membrane Biogenesis. Annu. Rev. Microbiol. 71, 539–556 (2017).

3. R. Miyazaki, M. Ai, N. Tanaka, T. Suzuki, N. Dhomae, T. Tsukazaki, Y. Akiyama, H. Mori, Inner membrane YfgM–PpiD heterodimer acts as a functional unit that associates with the SecY/E/G translocon and promotes protein translocation. J. Biol. Chem. 298 (2022).

4. W. J. Allen, S. L. Williams, I. Collinson, The Great Escape: Protein Trafficking from the Bacterial Cytosol to the Outer Membrane, Annual Review of Biochemistry. 95 (2026)pp. 431–454.

5. M. T. Doyle, H. D. Bernstein, Molecular Machines that Facilitate Bacterial Outer Membrane Protein Biogenesis, Annual Review of Biochemistry. 93 (2024)pp. 211–231.

6. T. Tsukazaki, H. Mori, Y. Echizen, R. Ishitani, S. Fukai, T. Tanaka, A. Perederina, D. G. Vassylyev, T. Kohno, A. D. Maturana, K. Ito, O. Nureki, Structure and function of a protein export-enhancing membrane component SecDF. Nature, doi: 10.1038/nature09980 (2011).

7. A. N. Combs, T. J. Silhavy, Periplasmic Chaperones: Outer Membrane Biogenesis and Envelope Stress. Annu. Rev. Microbiol. 78, 191–211 (2024).

8. T. Devlin, K. G. Fleming, A team of chaperones play to win in the bacterial periplasm. Trends Biochem. Sci. 49, 667–680 (2024).

9. Y. Gu, H. Li, H. Dong, Y. Zeng, Z. Zhang, N. G. Paterson, P. J. Stansfeld, Z. Wang, Y. Zhang, W. Wang, C. Dong, Structural basis of outer membrane protein insertion by the BAM complex. Nature 531, 64–69 (2016).

10. J. Bakelar, S. K. Buchanan, N. Noinaj, The structure of the β-barrel assembly machinery complex. Science 351, 180–186 (2016).

11. K. L. Fenn, J. E. Horne, J. A. Crossley, N. Böhringer, R. J. Horne, T. F. Schäberle, A. N. Calabrese, S. E. Radford, N. A. Ranson, Outer membrane protein assembly mediated by BAM-SurA complexes. Nat. Commun. 15, 7612–7612 (2024).

12. P. A. Lehner, M. Degen, R. P. Jakob, S. M. Modaresi, M. Callon, B. M. Burmann, T. Maier, S. Hiller, Architecture and conformational dynamics of the BAM-SurA holo insertase complex. Sci. Adv. 11 (2025).

13. R. Miyazaki, H. Kohga, N. Matsuoka, Y. Maruno, W. Yoshimoto, Y. S. Takahashi, D. H. Y. Yanto, Y. Nugraha, H. Shigematsu, T. Shiota, T. Tsukazaki, Cryo-EM structures of the BAM–P1/P2-visible SurA complex reveal dynamic and cooperative interactions in outer membrane protein assembly. bioRxiv, 2025.11.10.687524 (2026).

14. E. M. Germany, N. Thewasano, K. Imai, Y. Maruno, R. S. Bamert, C. J. Stubenrauch, R. A. Dunstan, Y. Ding, Y. Nakajima, X. Lai, C. T. Webb, K. Hidaka, K. S. Tan, H. Shen, T. Lithgow, T. Shiota, Dual recognition of multiple signals in bacterial outer membrane proteins enhances assembly and maintains membrane integrity. eLife 12, RP90274–RP90274 (2024).

15. D. Tomasek, S. Rawson, J. Lee, J. S. Wzorek, S. C. Harrison, Z. Li, D. Kahne, Structure of a nascent membrane protein as it folds on the BAM complex. Nature 583, 473–478 (2020).

16. R. Wu, J. W. Bakelar, K. Lundquist, Z. Zhang, K. M. Kuo, D. Ryoo, Y. T. Pang, C. Sun, T. White, T. Klose, W. Jiang, J. C. Gumbart, N. Noinaj, Plasticity within the barrel domain of BamA mediates a hybrid-barrel mechanism by BAM. Nat. Commun. 12, 7131–7131 (2021).

17. M. T. Doyle, J. R. Jimah, T. Dowdy, S. I. Ohlemacher, M. Larion, J. E. Hinshaw, H. D. Bernstein, Cryo-EM structures reveal multiple stages of bacterial outer membrane protein folding. Cell 185, 1143–1156.e13 (2022).

18. C. Shen, S. Chang, Q. Luo, K. C. Chan, Z. Zhang, B. Luo, T. Xie, G. Lu, X. Zhu, X. Wei, C. Dong, R. Zhou, X. Zhang, X. Tang, H. Dong, Structural basis of BAM-mediated outer membrane β-barrel protein assembly. Nature 617, 185–193 (2023).

19. B. D. Thomson, M. D. Marquez, S. Rawson, T. M. A. dos Santos, S. C. Harrison, D. Kahne, Structures of folding intermediates on BAM show diverse substrates fold by a conserved mechanism. Proc. Natl. Acad. Sci. 123, e2534936123 (2026).

20. S. Okuda, D. J. Sherman, T. J. Silhavy, N. Ruiz, D. Kahne, Lipopolysaccharide transport and assembly at the outer membrane: the PEZ model. Nat. Rev. Microbiol. 14, 337–345 (2016).

21. L. Törk, C. B. Moffatt, T. G. Bernhardt, E. C. Garner, D. Kahne, Single-molecule dynamics show a transient lipopolysaccharide transport bridge. Nature 623, 814–819 (2023).

22. S. Qiao, Q. Luo, Y. Zhao, X. C. Zhang, Y. Huang, Structural basis for lipopolysaccharide insertion in the bacterial outer membrane. Nature 511, 108–111 (2014).

23. H. Dong, Q. Xiang, Y. Gu, Z. Wang, N. G. Paterson, P. J. Stansfeld, C. He, Y. Zhang, W. Wang, C. Dong, Structural basis for outer membrane lipopolysaccharide insertion. Nature 511, 52–56 (2014).

24. S. Narita, C. Masui, T. Suzuki, N. Dohmae, Y. Akiyama, Protease homolog BepA (YfgC) promotes assembly and degradation of β-barrel membrane proteins in Escherichia coli. Proc. Natl. Acad. Sci. U. S. A. 110, E3612–21 (2013).

25. S.-S. Chng, M. Xue, R. A. Garner, H. Kadokura, D. Boyd, J. Beckwith, D. Kahne, Disulfide Rearrangement Triggered by Translocon Assembly Controls Lipopolysaccharide Export. Science 337, 1665–1668 (2012).

26. J. Lee, M. Xue, J. S. Wzorek, T. Wu, M. Grabowicz, L. S. Gronenberg, H. A. Sutterlin, R. M. Davis, N. Ruiz, T. J. Silhavy, D. E. Kahne, Characterization of a stalled complex on the β-barrel assembly machine. Proc. Natl. Acad. Sci. 113, 8717–8722 (2016).

27. Y. Yang, H. Chen, R. A. Corey, V. Morales, Y. Quentin, C. Froment, A. Caumont-Sarcos, C. Albenne, O. Burlet-Schiltz, D. Ranava, P. J. Stansfeld, J. Marcoux, R. Ieva, LptM promotes oxidative maturation of the lipopolysaccharide translocon by substrate binding mimicry. Nat. Commun. 14, 6368–6368 (2023).

28. R. Miyazaki, M. Kimoto, H. Kohga, T. Tsukazaki, Structural basis of lipopolysaccharide translocon assembly mediated by the small lipoprotein LptM. Cell Rep. 44, 116013 (2025).

29. G. R. Soltes, N. R. Martin, E. Park, H. A. Sutterlin, T. J. Silhavy, Distinctive roles for periplasmic proteases in the maintenance of essential outer membrane protein assembly. J. Bacteriol. 199, e00418–17 (2017).

30. Y. Daimon, C. Iwama-Masui, Y. Tanaka, T. Shiota, T. Suzuki, R. Miyazaki, H. Sakurada, T. Lithgow, N. Dohmae, H. Mori, T. Tsukazaki, S.-I. Narita, Y. Akiyama, The TPR domain of BepA is required for productive interaction with substrate proteins and the β-barrel assembly machinery complex. Mol. Microbiol. 106, 760–776 (2017).

31. R. Miyazaki, T. Watanabe, K. Yoshitani, Y. Akiyama, Edge-strand of BepA interacts with immature LptD on the β-barrel assembly machine to direct it to on- and off-pathways. eLife 10 (2021).

32. M. Shahrizal, Y. Daimon, Y. Tanaka, Y. Hayashi, S. Nakayama, S. Iwaki, S. Narita, H. Kamikubo, Y. Akiyama, T. Tsukazaki, Structural Basis for the Function of the β-Barrel Assembly-Enhancing Protease BepA. J. Mol. Biol. 431, 625–635 (2019).

33. Y. Daimon, S.-I. Narita, R. Miyazaki, Y. Hizukuri, H. Mori, Y. Tanaka, T. Tsukazaki, Y. Akiyama, Reversible autoinhibitory regulation of Escherichia coli metallopeptidase BepA for selective β-barrel protein degradation. Proc. Natl. Acad. Sci. U. S. A. 117, 27989–27996 (2020).

34. J. A. Bryant, I. T. Cadby, Z.-S. Chong, G. Boelter, Y. R. Sevastsyanovich, F. C. Morris, A. F. Cunningham, G. Kritikos, R. W. Meek, M. Banzhaf, S.-S. Chng, A. L. Lovering, I. R. Henderson, Structure-Function Characterization of the Conserved Regulatory Mechanism of the Escherichia coli M48 Metalloprotease BepA. J. Bacteriol. 203 (2020).

35. M. Gao, D. Nakajima An, J. Skolnick, Deep learning-driven insights into super protein complexes for outer membrane protein biogenesis in bacteria. eLife 11, e82885 (2022).

36. Y. Imaizumi, K. Takanuki, T. Miyake, M. Takemoto, K. Hirata, M. Hirose, R. Oi, T. Kobayashi, K. Miyoshi, R. Aruga, T. Yokoyama, S. Katagiri, H. Matsuura, K. Iwasaki, T. Kato, M. K. Kaneko, Y. Kato, M. Tajiri, S. Akashi, O. Nureki, Y. Hizukuri, Y. Akiyama, T. Nogi, Mechanistic insights into intramembrane proteolysis by E. coli site-2 protease homolog RseP. Sci. Adv. 8, eabp9011.

37. K. L. Fenn, V. Higgins, J. M. Machin, A. N. Calabrese, A. Berry, S. E. Radford, N. A. Ranson, BAM-BepA complexes in outer membrane protein quality control. Nat. Commun., doi: 10.1038/s41467-026-75227-x (2026).

38. H. Voedts, P. C. Nguyen, V.-S. Nguyen, P. Leverrier, B. Iorga, S. H. Cho, H. Remaut, J.-F. Collet, Stress-induced Membrane Insertion at the β-Barrel Assembly Machinery Complex Regulates BepA Metalloprotease Activity. bioRxiv, 2026.07.01.735768 (2026).

39. Y. Imai, K. J. Meyer, A. Iinishi, Q. Favre-Godal, R. Green, S. Manuse, M. Caboni, M. Mori, S. Niles, M. Ghiglieri, C. Honrao, X. Ma, J. J. Guo, A. Makriyannis, L. Linares-Otoya, N. Böhringer, Z. G. Wuisan, H. Kaur, R. Wu, A. Mateus, A. Typas, M. M. Savitski, J. L. Espinoza, A. O’Rourke, K. E. Nelson, S. Hiller, N. Noinaj, T. F. Schäberle, A. D’Onofrio, K. Lewis, A new antibiotic selectively kills Gram-negative pathogens. Nature 576, 459–464 (2019).

40. H. Kaur, R. P. Jakob, J. K. Marzinek, R. Green, Y. Imai, J. R. Bolla, E. Agustoni, C. V. Robinson, P. J. Bond, K. Lewis, T. Maier, S. Hiller, The antibiotic darobactin mimics a β-strand to inhibit outer membrane insertase. Nature 593, 125–129 (2021).

41. K. M. Storek, D. Sun, S. T. Rutherford, Inhibitors targeting BamA in gram-negative bacteria. Biochim. Biophys. Acta BBA - Mol. Cell Res. 1871, 119609 (2024).

42. F. Munder, M. D. Johnson, I. Samuels, L. McCaughey, O. Zdorevskyi, C. Wang, A. Kropp, L. Zavan, E. P. Price, D. S. Sarovich, S. Varshney, C. A. McDevitt, H. Venugopal, V. Sharma, M. T. Doyle, F. Short, D. Ghosal, J. P. R. Connolly, G. J. Knott, R. Grinter, L-type pyocins inhibit the BAM complex to kill without cell entry. Nat. Commun., doi: 10.1038/s41467-026-74995-w (2026).

43. T. Baba, T. Ara, M. Hasegawa, Y. Takai, Y. Okumura, M. Baba, K. A. Datsenko, M. Tomita, B. L. Wanner, H. Mori, Construction of Escherichia coli K-12 in-frame, single-gene knockout mutants: the Keio collection. Mol. Syst. Biol. 2, 2006.0008-2006.0008 (2006).

44. H. Mori, K. Ito, The long alpha-helix of SecA is important for the ATPase coupling of translocation. J. Biol. Chem. 281, 36249–36256 (2006).

45. K. Kanehara, K. Ito, Y. Akiyama, YaeL proteolysis of RseA is controlled by the PDZ domain of YaeL and a Gln-rich region of RseA. EMBO J. 22, 6389–6398 (2003).

46. S. D. Gunasinghe, T. Shiota, C. J. Stubenrauch, K. E. Schulze, C. T. Webb, A. J. Fulcher, R. A. Dunstan, I. D. Hay, T. Naderer, D. R. Whelan, T. D. M. Bell, K. D. Elgass, R. A. Strugnell, T. Lithgow, The WD40 Protein BamB Mediates Coupling of BAM Complexes into Assembly Precincts in the Bacterial Outer Membrane. Cell Rep. 23, 2782–2794 (2018).

47. A. Kihara, Y. Akiyama, K. Ito, A protease complex in the Escherichia coli plasma membrane: HflKC (HflA) forms a complex with FtsH (HflB), regulating its proteolytic activity against SecY. EMBO J. 15, 6122–6131 (1996).

48. T. Baba, A. Jacq, E. Brickman, J. Beckwith, T. Taura, C. Ueguchi, Y. Akiyama, K. Ito, Characterization of cold-sensitive secY mutants of Escherichia coli. J. Bacteriol. 172, 7005–7010 (1990).

49. Y. Sugano, A. Furukawa, O. Nureki, Y. Tanaka, T. Tsukazaki, SecY-SecA fusion protein retains the ability to mediate protein transport. PLOS ONE 12, e0183434–e0183434 (2017).

50. D. N. Mastronarde, Automated electron microscope tomography using robust prediction of specimen movements. J. Struct. Biol. 152, 36–51 (2005).

51. A. Punjani, J. L. Rubinstein, D. J. Fleet, M. A. Brubaker, cryoSPARC: algorithms for rapid unsupervised cryo-EM structure determination. Nat. Methods 14, 290–296 (2017).

52. J. Jumper, R. Evans, A. Pritzel, T. Green, M. Figurnov, O. Ronneberger, K. Tunyasuvunakool, R. Bates, A. Žídek, A. Potapenko, A. Bridgland, C. Meyer, S. A. A. Kohl, A. J. Ballard, A. Cowie, B. Romera-Paredes, S. Nikolov, R. Jain, J. Adler, T. Back, S. Petersen, D. Reiman, E. Clancy, M. Zielinski, M. Steinegger, M. Pacholska, T. Berghammer, S. Bodenstein, D. Silver, O. Vinyals, A. W. Senior, K. Kavukcuoglu, P. Kohli, D. Hassabis, Highly accurate protein structure prediction with AlphaFold. Nature 596, 583–589 (2021).

53. J. Abramson, J. Adler, J. Dunger, R. Evans, T. Green, A. Pritzel, O. Ronneberger, L. Willmore, A. J. Ballard, J. Bambrick, S. W. Bodenstein, D. A. Evans, C.-C. Hung, M. O’Neill, D. Reiman, K. Tunyasuvunakool, Z. Wu, A. Žemgulytė, E. Arvaniti, C. Beattie, O. Bertolli, A. Bridgland, A. Cherepanov, M. Congreve, A. I. Cowen-Rivers, A. Cowie, M. Figurnov, F. B. Fuchs, H. Gladman, R. Jain, Y. A. Khan, C. M. R. Low, K. Perlin, A. Potapenko, P. Savy, S. Singh, A. Stecula, A. Thillaisundaram, C. Tong, S. Yakneen, E. D. Zhong, M. Zielinski, A. Žídek, V. Bapst, P. Kohli, M. Jaderberg, D. Hassabis, J. M. Jumper, Accurate structure prediction of biomolecular interactions with AlphaFold 3. Nature 630, 493–500 (2024).

54. E. F. Pettersen, T. D. Goddard, C. C. Huang, E. C. Meng, G. S. Couch, T. I. Croll, J. H. Morris, T. E. Ferrin, UCSF ChimeraX: Structure visualization for researchers, educators, and developers. Protein Sci. Publ. Protein Soc. 30, 70–82 (2021).

55. P. Emsley, B. Lohkamp, W. G. Scott, K. Cowtan, Features and development of Coot. Acta Crystallogr. D Biol. Crystallogr. 66, 486–501 (2010).

56. P. D. Adams, P. V. Afonine, G. Bunkóczi, V. B. Chen, I. W. Davis, N. Echols, J. J. Headd, L.-W. Hung, G. J. Kapral, R. W. Grosse-Kunstleve, A. J. McCoy, N. W. Moriarty, R. Oeffner, R. J. Read, D. C. Richardson, J. S. Richardson, T. C. Terwilliger, P. H. Zwart, PHENIX: a comprehensive Python-based system for macromolecular structure solution. Acta Crystallogr. D Biol. Crystallogr. 66, 213–221 (2010).

57. J. W. Chin, P. G. Schultz, In vivo photocrosslinking with unnatural amino Acid mutagenesis. *Chembiochem Eur*. J. Chem. Biol. 3, 1135–1137 (2002).

58. T. S. Young, I. Ahmad, J. A. Yin, P. G. Schultz, An enhanced system for unnatural amino acid mutagenesis in E. coli. J. Mol. Biol. 395, 361–374 (2010).

59. R. Miyazaki, Y. Akiyama, H. Mori, A photo-cross-linking approach to monitor protein dynamics in living cells. Biochim. Biophys. Acta Gen. Subj. 1864, 129317–129317 (2020).

60. A. Kihara, Y. Akiyama, K. Ito, FtsH is required for proteolytic elimination of uncomplexed forms of SecY, an essential protein translocase subunit. Proc. Natl. Acad. Sci. U. S. A. 92, 4532–4536 (1995).

61. T. J. Silhavy, M. L. Berman, L. W. (Lynn W.) Enquist, C. S. H. Laboratory., Experiments with Gene Fusions (Cold Spring Harbor Laboratory, Cold Spring Harbor, N.Y, 1984).

62. Y. Akiyama, T. Ogura, K. Ito, Involvement of FtsH in protein assembly into and through the membrane. I. Mutations that reduce retention efficiency of a cytoplasmic reporter. J. Biol. Chem. 269, 5218–5224 (1994).

63. G. Roman-Hernandez, J. H. Peterson, H. D. Bernstein, Reconstitution of bacterial autotransporter assembly using purified components. eLife 3, e04234 (2014).

