## Supplementary Materials for "Structural and functional insights into multiple BAM-bound conformations of BepA enabling substrate triage at the outer membrane"

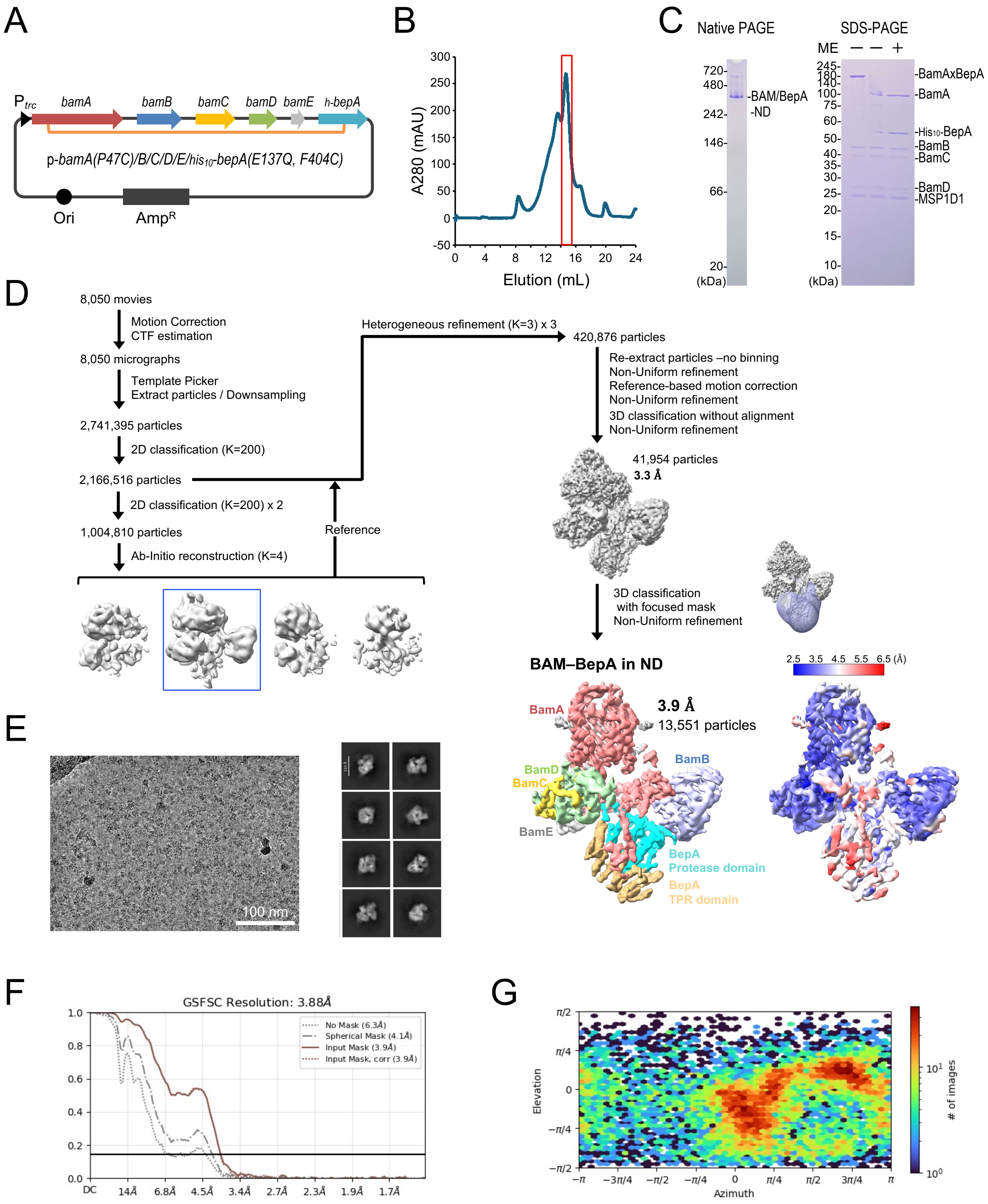


**fig. S1. Cryo-EM analysis of the nanodisc-reconstituted BAM–BepA complex.**
(**A**) Schematic of a plasmid encoding *bamA(P47C)/B/C/D/E/his$_{10}$-bepA(E137Q, F404C)*. N-terminal His$_{10}$-tag is inserted after the signal sequence of BepA. (**B**) Size-exclusion chromatogram of the nanodisc-reconstituted BAM–BepA complex using a Superose 6 Increase column. (**C**) Native PAGE and SDS–PAGE analyses of the nanodisc-reconstituted BAM–BepA complex stained with CBB. (**D**) Cryo-EM data-processing workflow for the BAM–BepA complex. Final cryo-EM maps of the nanodisc-reconstituted BAM–BepA complex. The map is displayed at contour level 0.03. BepA TPR and protease domains are colored in orange and cyan, respectively. BamA, BamB, BamC, BamD, and BamE are in red, light blue, yellow, light green, and gray, respectively. The local resolution of the cryo-EM map was calculated using CryoSPARC and is shown. (**E**) Representative cryo-EM micrograph and 2D classes. (**F**) Gold-standard Fourier shell correlation (GSFSC) curve used for global-resolution estimates within CryoSPARC. (**G**) Angular distribution of particles used for the final 3D reconstruction.

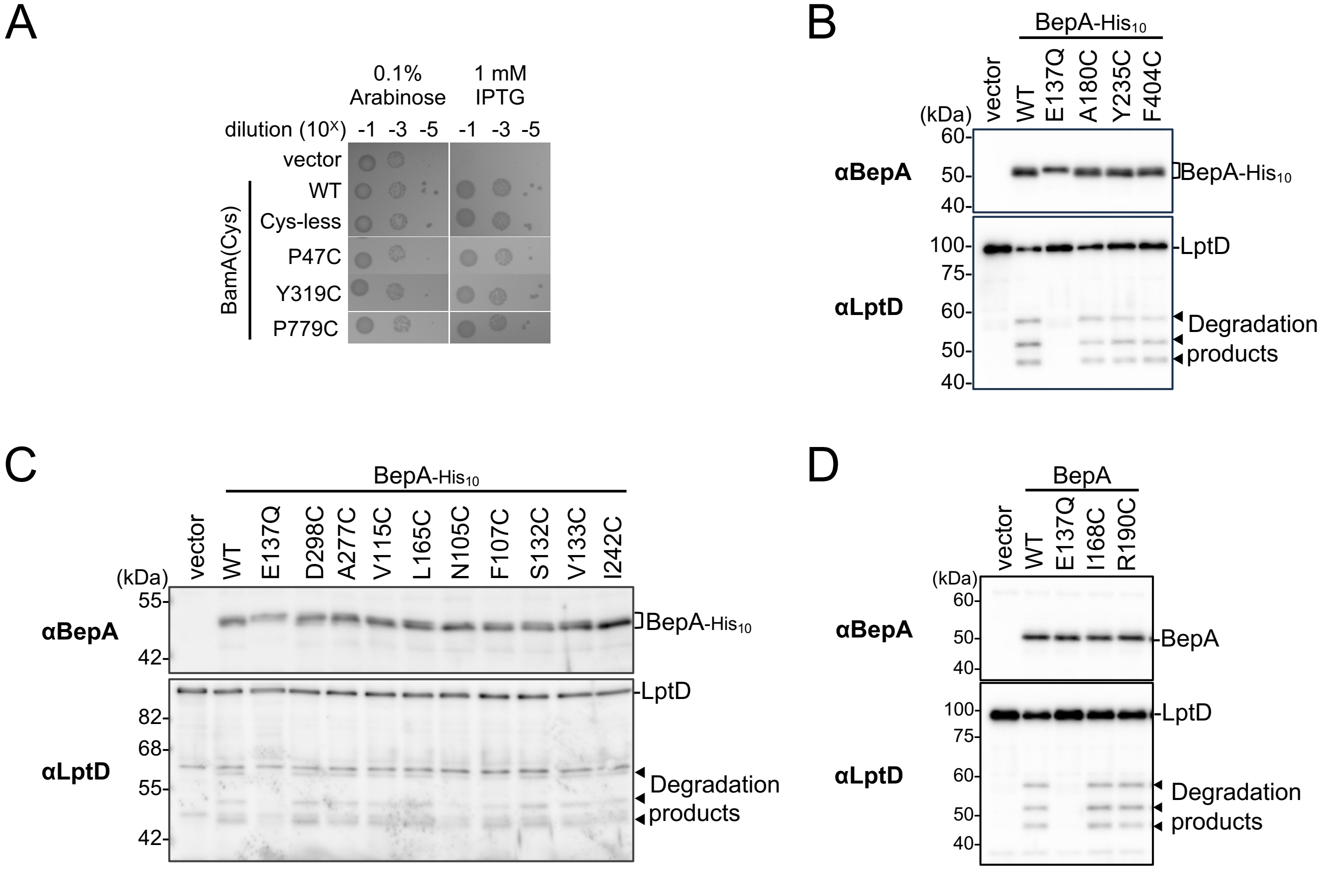


**fig. S2. Activities of BamA and BepA cysteine derivatives used in this study.**

(**A**) Complementation activities of the BamA Cys derivatives. Cells of RM4278 (P$_{ara}$-*bamA*) carrying pEVOL-pBpF and either pUC118 or pUC118-*bamA(Cys)* plasmid were grown at 30 ºC in L-medium supplemented with 0.1% arabinose for 2.5 h. Cells were washed, resuspended in saline, and serially diluted with saline (to about $10^9$ cells/mL). Aliquots (2.5 µL) of each dilution were spotted onto LB-agar plates containing 0.1% arabinose (positive control) or 1 mM IPTG (for the expression of the BamA Cys mutants from a plasmid). Plates were incubated at 30 ºC for 22 h. (**B–D**) Protease activities of the BepA Cys mutants against overproduced LptD. (B) Cells of SN56 (Δ*bepA*) carrying pTWV228-*lptD-his$_{10}$* and either pSTD689 or pSTD689-*bepA(mut)-his$_{10}$* plasmids were grown at 30 ºC in LB-medium until early log phase and induced with 1 mM IPTG for 1 h. Total cellular proteins were acid-precipitated and analyzed by 7.5 or 10% Laemmli SDS-PAGE and immunoblotting with the indicated antibodies. (C) Cells of SN56 (Δ*bepA*) carrying pTWV228-*lptD-his$_{10}$* and either pHSG575 or pHSG575-*bepA(mut)-his$_{10}$* plasmids were grown, induced, and analyzed as in (B). (D) Cells of SN56 (Δ*bepA*) carrying pTWV228-*lptD-his$_{10}$* and either pSTD689 or pSTD689-*bepA(mut)* plasmids were grown, induced, and analyzed as in (B). The C-terminal His$_{10}$-tag of BepA is self-cleaved *in vivo*[24]. The slower migration of the E137Q mutant in **b** and **c** likely reflects reduced self-cleavage of the C-terminal His$_{10}$-tag.

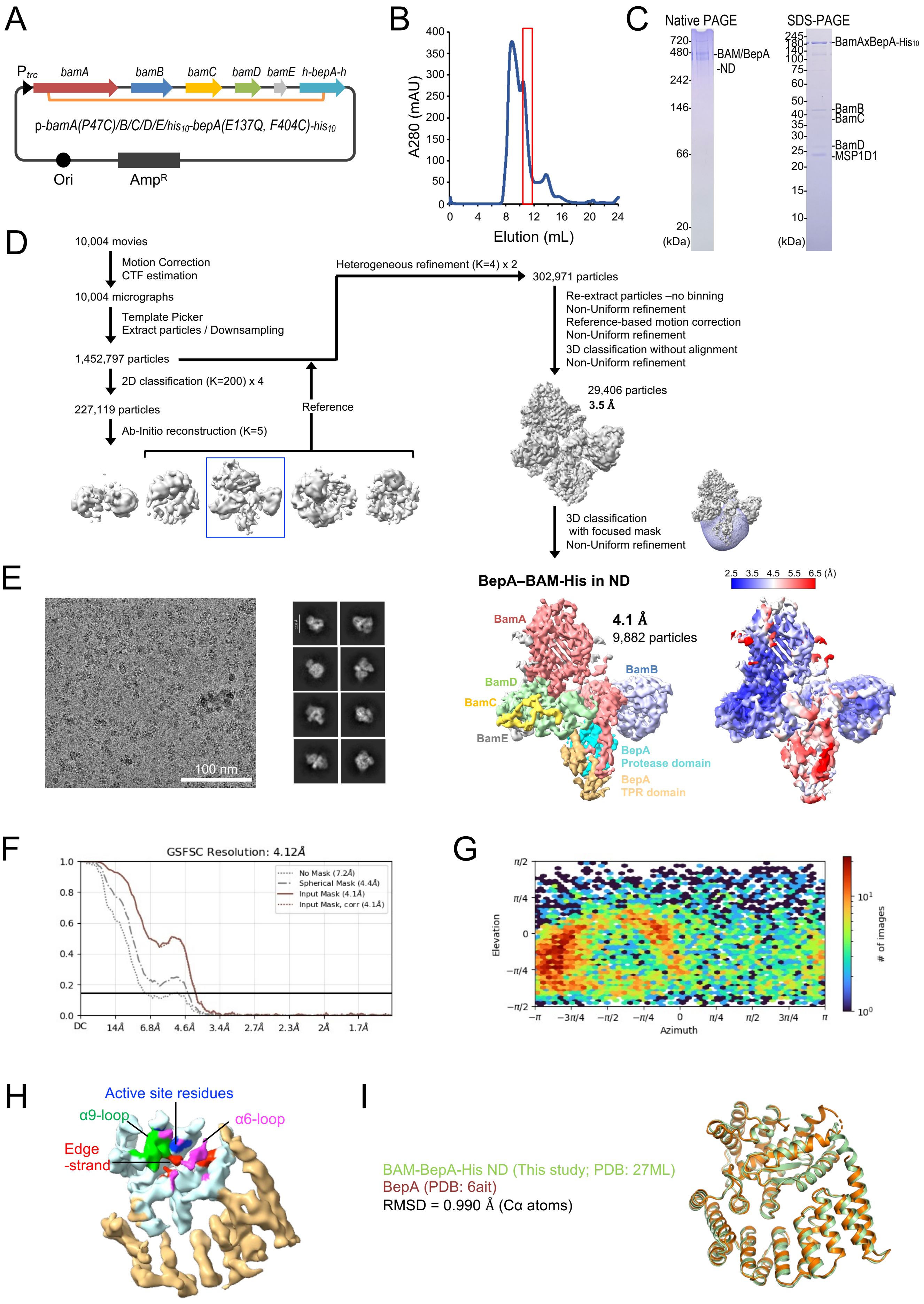
A
P trc
bamA
bamB
bamC
bamD
bamE
h-bepA-h
p-bamA(P47C)/B/C/D/E/his10-bepA(E137Q, F404C)-his10
Ori
AmpR
B
A280 (mAU)
Elution (mL)
C
Native PAGE
720-
480-
242-
146-
66-
20-
(kDa)
-BAM/BepA
-ND
SDS-PAGE
245-
180-
140-
100-
75-
60-
50-
40-
35-
30-
25-
20-
15-
10-
(kDa)
-BamAxBepA-His10
-BamB
-BamC
-BamD
-MSP1D1
D
10,004 movies
Motion Correction
CTF estimation
10,004 micrographs
Template Picker
Extract particles / Downsampling
1,452,797 particles
2D classification (K=200) x 4
227,119 particles
Ab-Initio reconstruction (K=5)
Reference
Heterogeneous refinement (K=4) x 2
302,971 particles
Re-extract particles –no binning
Non-Uniform refinement
Reference-based motion correction
Non-Uniform refinement
3D classification without alignment
Non-Uniform refinement
29,406 particles
3.5 Å
3D classification
with focused mask
Non-Uniform refinement
E
100 nm
BepA–BAM-His in ND
2.5 3.5 4.5 5.5 6.5 (Å)
BamA
4.1 Å
9,882 particles
BamB
BamD
BamC
BamE
BepA
Protease domain
BepA
TPR domain
F
GSFSC Resolution: 4.12Å
No Mask (7.2Å)
Spherical Mask (4.4Å)
Input Mask (4.1Å)
Input Mask, corr (4.1Å)
DC
14Å
6.8Å
4.6Å
3.4Å
2.7Å
2.3Å
2Å
1.7Å
G
Elevation
Azimuth
### of images
H
Active site residues
α9-loop
α6-loop
Edge
-strand
I
BAM-BepA-His ND (This study; PDB: 27ML)
BepA (PDB: 6ait)
RMSD = 0.990 Å (Cα atoms)

**fig. S3. Cryo-EM analysis of the nanodisc-reconstituted BAM–BepA-His complex.**

(**A**) Schematic of a plasmid encoding *bamA(P47C)/B/C/D/E/his$_{10}$-bepA(E137Q, F404C)-his$_{10}$*. N-terminal His$_{10}$-tag is inserted after the signal sequence of BepA. (**B**) Size-exclusion chromatogram of the nanodisc-reconstituted BAM–BepA-His complex using a Superdex 200 Increase column. (**C**) Native PAGE and SDS–PAGE analyses of the nanodisc-reconstituted BAM–BepA-His complex stained with CBB. (**D**) Cryo-EM data-processing workflow for the BAM–BepA-His complex. Final cryo-EM maps of the nanodisc-reconstituted BAM–BepA-His complex. The map is displayed at contour level 0.03. BepA TPR and protease domains are colored in orange and cyan, respectively. BamA, BamB, BamC, BamD, and BamE are in red, light blue, yellow, light green, and gray, respectively. The local resolution of the cryo-EM map was calculated using CryoSPARC and is shown. (**E**) Representative cryo-EM micrograph and 2D classes. (**F**) Gold-standard Fourier shell correlation (GSFSC) curve used for global-resolution estimates within CryoSPARC. (**G**) Angular distribution of particles used for the final 3D reconstruction. (**H**) Cryo-EM map of BepA in in the BAM–BepA complex shown in (D). The protease and the TPR domains of BepA are shown in light cyan and orange, respectively. The edge-strand, the proteolytic active-site residues (the HExxH motif and the third zinc ligand, Glu-201), the α6-loop, and the α9-loop in the protease domain are shown in red, blue, magenta, and green, respectively. (**I**) Structural comparison of BepA in the nanodisc-reconstituted BAM–BepA-His complex with the BepA crystal structure. BepA in the BAM–BepA-His complex and the BepA crystal structure are shown in light green and orange, respectively.

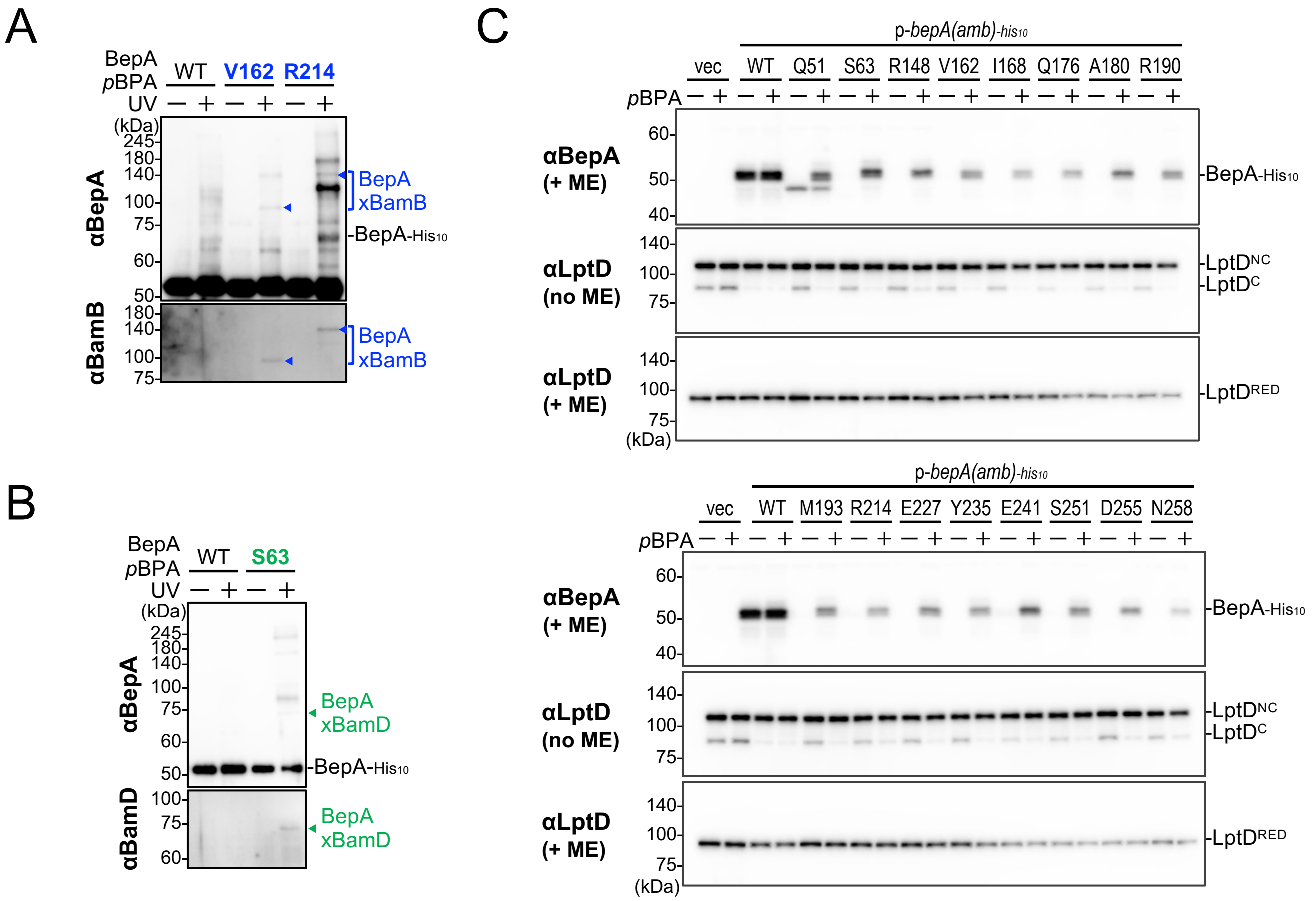


**fig. S4.** ***In vivo*** **photo-crosslinking analysis using BepA(*p*BPA) derivatives.**
(**A, B**) *In vivo* photo-crosslinking analysis of BepA. The same samples in Fig. 2A were analyzed by SDS–PAGE followed by immunoblotting using the indicated antibodies. (**C**) Accumulation of the BepA(*p*BPA) derivatives and $LptD^{C}$ in the Δ*bepA* cells expressing the BepA derivatives. Cells of SN56 (Δ*bepA*) carrying pEVOL-pBpF and either pUC18 or pUC18-*bepA(amb)-his*$_{10}$ plasmids were grown at 30 ºC in LB-medium until early log phase and induced with 1 mM IPTG for 1 h. Total cellular proteins were acid-precipitated, and analyzed by 7.5 or 10% Laemmli SDS-PAGE under a reducing (+ME) or non-reducing (no ME) conditions and immunoblotting with indicated antibodies.

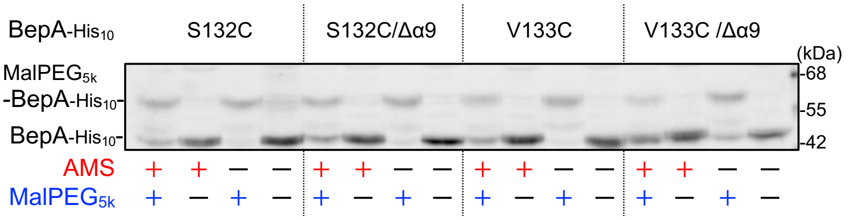


**fig. S5. Effect of α9-loop deletion on the AMS accessibility of the BepA S132C and V133C derivatives.** Cells of SN56 (Δ*bepA*) carrying pHSG575-*bepA(Cys, mut)-his₁₀* were grown and analyzed as in Fig. 2E.

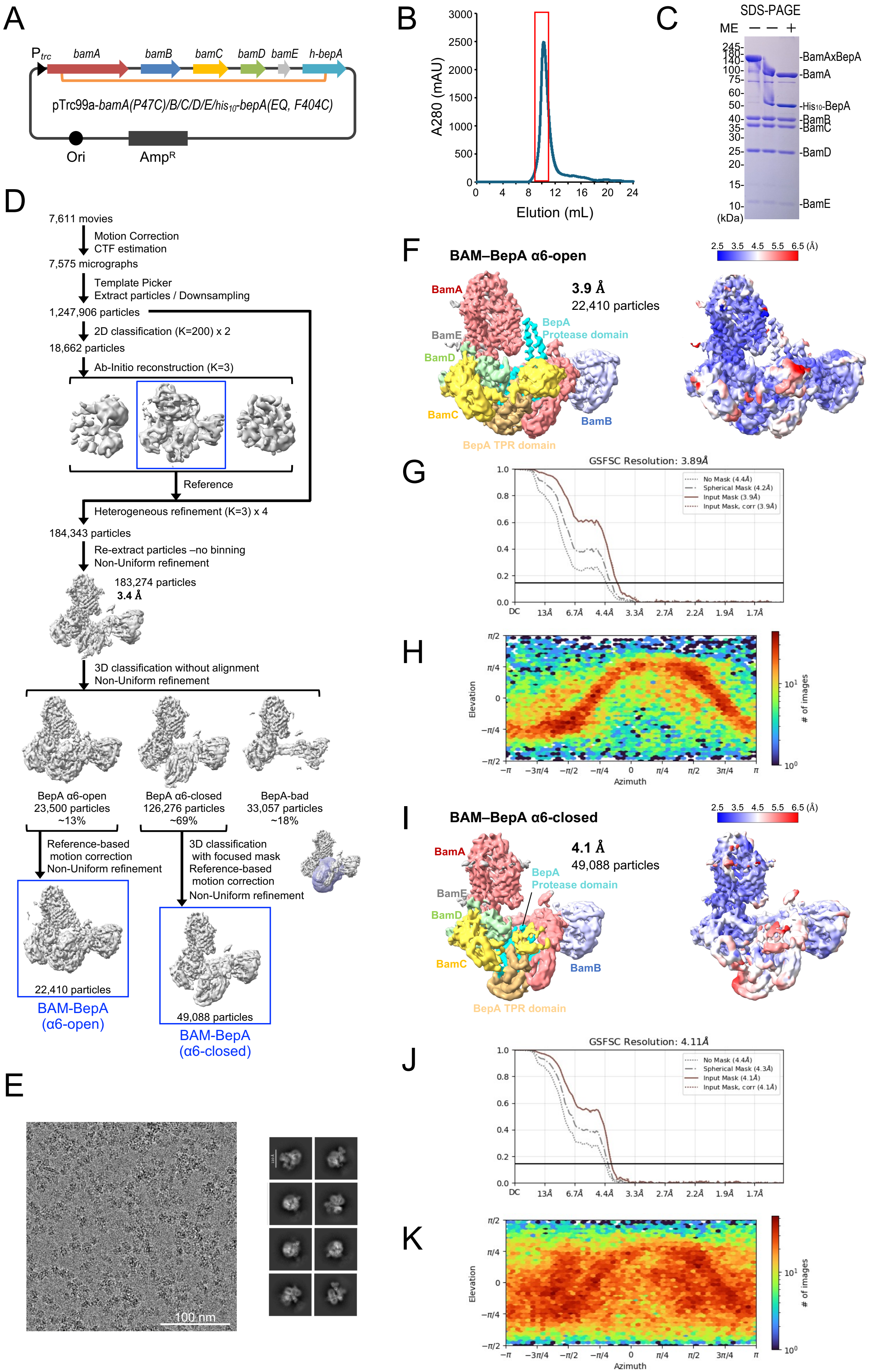

A
Ptrc
bamA
bamB
bamC
bamD
bamE
h-bepA
pTrc99a-bamA(P47C)/B/C/D/E/his10-bepA(EQ, F404C)
Ori
AmpR
B
A280 (mAU)
Elution (mL)
C
SDS-PAGE
ME
(kDa)
-BamAxBepA
-BamA
-His10-BepA
-BamB
-BamC
-BamD
-BamE
D
7,611 movies
Motion Correction
CTF estimation
7,575 micrographs
Template Picker
Extract particles / Downsampling
1,247,906 particles
2D classification (K=200) x 2
18,662 particles
Ab-Initio reconstruction (K=3)
Reference
Heterogeneous refinement (K=3) x 4
184,343 particles
Re-extract particles –no binning
Non-Uniform refinement
183,274 particles
3.4 Å
3D classification without alignment
Non-Uniform refinement
BepA α6-open
23,500 particles
~13%
BepA α6-closed
126,276 particles
~69%
BepA-bad
33,057 particles
~18%
Reference-based
motion correction
Non-Uniform refinement
3D classification
with focused mask
Reference-based
motion correction
Non-Uniform refinement
22,410 particles
BAM-BepA
(α6-open)
49,088 particles
BAM-BepA
(α6-closed)
E
100 nm
F
BAM–BepA α6-open
3.9 Å
22,410 particles
BamA
BamE
BamD
BamC
BepA
Protease domain
BamB
BepA TPR domain
2.5 3.5 4.5 5.5 6.5 (Å)
G
GSFSC Resolution: 3.89Å
No Mask (4.4Å)
Spherical Mask (4.2Å)
Input Mask (3.9Å)
Input Mask, corr (3.9Å)
H
Elevation
Azimuth
### of images
I
BAM–BepA α6-closed
4.1 Å
49,088 particles
BamA
BepA
Protease domain
BamE
BamD
BamC
BamB
BepA TPR domain
2.5 3.5 4.5 5.5 6.5 (Å)
J
GSFSC Resolution: 4.11Å
No Mask (4.4Å)
Spherical Mask (4.3Å)
Input Mask (4.1Å)
Input Mask, corr (4.1Å)
K
Elevation
Azimuth
### of images

**fig. S6. Cryo-EM analysis of the BAM–BepA complex.**
(**A**) Schematic of a plasmid encoding *bamA(P47C)/B/C/D/E/his$_{10}$-bepA(E137Q, F404C)*. N-terminal His$_{10}$-tag is inserted after the signal sequence of BepA. (**B**) Size-exclusion chromatogram of the purified BAM–BepA complex using a Superdex 200 Increase column. (**C**) SDS–PAGE analysis of the purified BAM–BepA complex stained with CBB. (**D**) Cryo-EM data-processing workflow for the BAM–BepA complex. (**E**) Representative cryo-EM micrograph and 2D classes. (**F**) Final cryo-EM maps of BAM–BepA α6-open. The map is displayed at contour level 0.0175. BepA TPR and protease domains are colored in orange and cyan, respectively. BamA, BamB, BamC, BamD, and BamE are in red, light blue, yellow, light green, and gray, respectively. The local resolution of the cryo-EM map was calculated using CryoSPARC and is shown. (**G**) Gold-standard Fourier shell correlation (GSFSC) curve used for global-resolution estimates within CryoSPARC. (**H**) Angular distribution of particles used for the final 3D reconstruction. (**I**) Final cryo-EM maps of BAM–BepA α6-closed. The map is displayed at contour level 0.016. The BepA domains and Bam factors are colored as in (F). The local resolution of the cryo-EM map was calculated using CryoSPARC and is shown. (**J**) Gold-standard Fourier shell correlation (GSFSC) curve used for global-resolution estimates within CryoSPARC. (**K**) Angular distribution of particles used for the final 3D reconstruction.

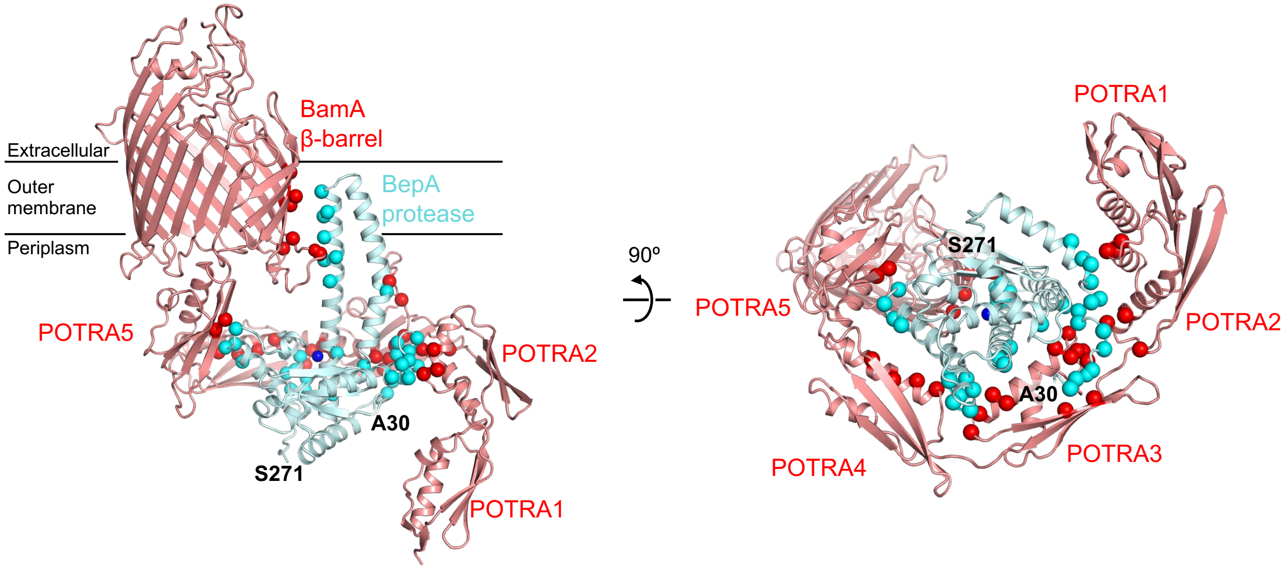


**fig. S7. Interface between BamA and the BepA protease domain in the α6-open/α9-closed BAM–BepA structure.**
BamA and the BepA protease domain are shown in red and pale cyan, respectively. BamA residues contacting BepA and BepA residues contacting BamA are shown as red and cyan spheres, respectively. The N- and C-terminal residues of the modeled BepA protease domain, A30 and S271, respectively, are labeled. BamB, BamC, BamD, BamE, and the BepA TPR domain are omitted for clarity.

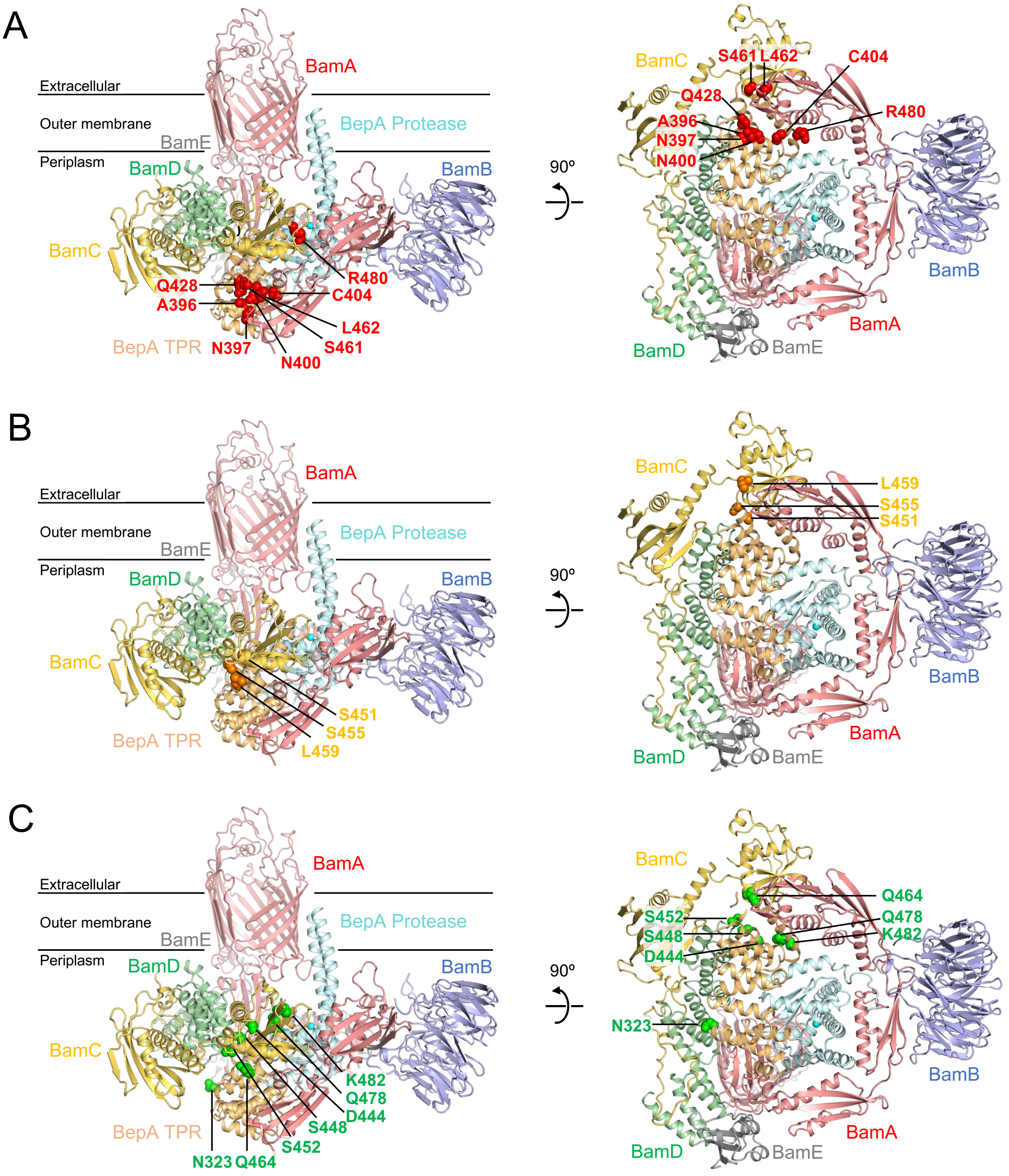


**fig. S8. Mapping of the BAM-crosslinking sites in the BepA TPR domain onto the BAM–BepA α6-open/α9-closed structure.**
(**A–C**) BepA residues crosslinked in the TPR domain to BamA (A), BamC (B), and BamD (C) are mapped onto the BAM–BepA α6-open/α9-closed structure. BamA, BamB, BamC, BamD, and BamE are shown in red, light blue, yellow, light green, and gray, respectively. The BepA protease and TPR domains are shown in pale cyan and orange, respectively. BepA residues crosslinked with BamA, BamC, and BamD are indicated by red (A), orange (B), and green (C) spheres, respectively.

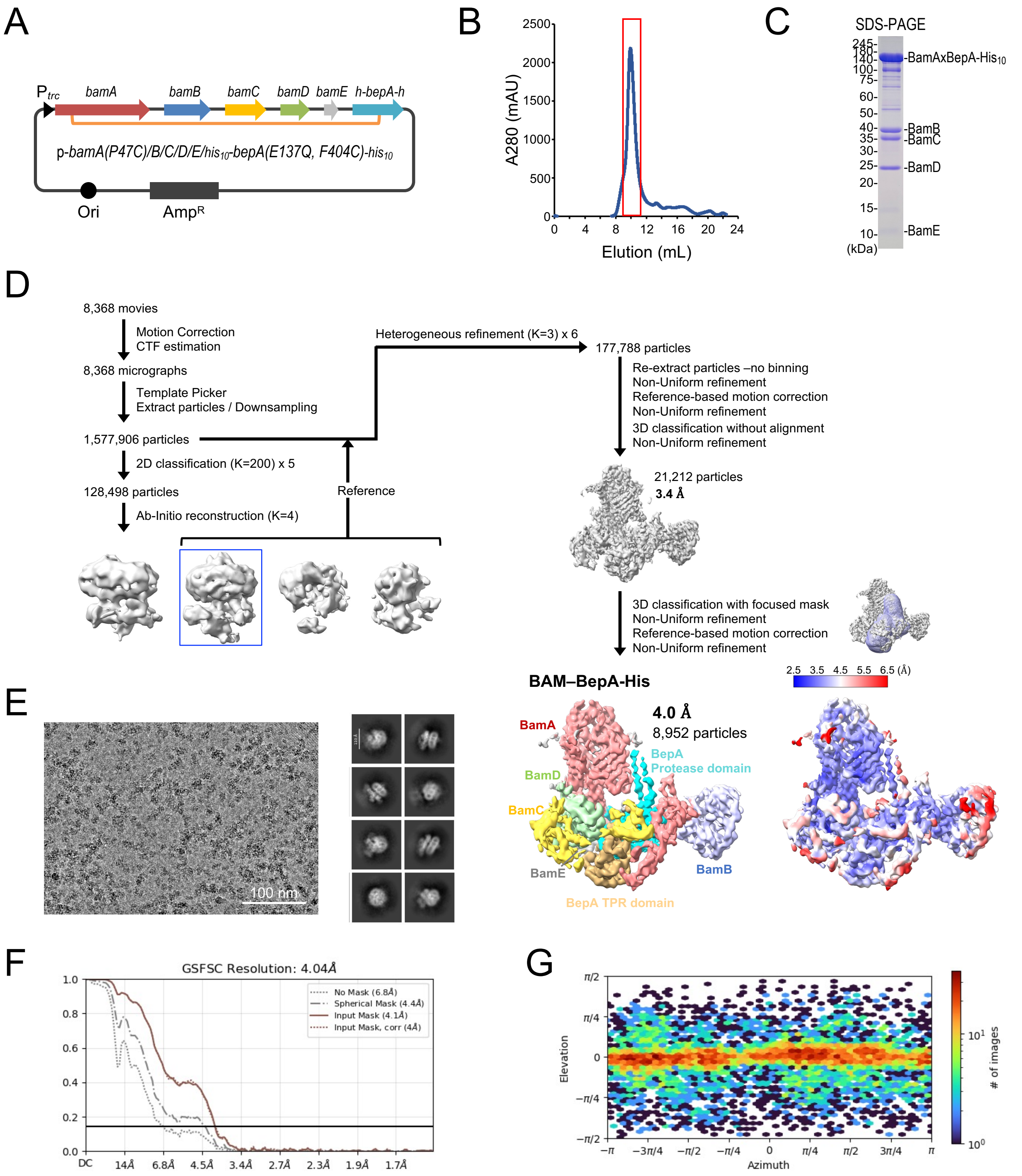


**fig. S9. Cryo-EM analysis of the BAM–BepA-His complex.**
(**A**) Schematic of a plasmid encoding *bamA(P47C)/B/C/D/E/his$_{10}$-bepA(E137Q, F404C)-his$_{10}$*. N-terminal His$_{10}$-tag is inserted after the signal sequence of BepA. (**B**) Size-exclusion chromatogram of the purified BAM–BepA-His complex using a Superdex 200 Increase column. (**C**) SDS–PAGE analysis of the purified BAM–BepA-His complex stained with CBB. (**D**) Cryo-EM data-processing workflow for the BAM–BepA-His complex. Final cryo-EM maps of the BAM–BepA-His complex. The map is displayed at contour level 0.028. BepA TPR and Protease domains are colored in orange and cyan, respectively. BamA, BamB, BamC, BamD, and BamE are in red, light blue, yellow, light green, and gray, respectively. The local resolution of the cryo-EM map was calculated using CryoSPARC and is shown. (**E**) Representative cryo-EM micrograph and 2D classes. (**F**) Gold-standard Fourier shell correlation (GSFSC) curve used for global-resolution estimates within CryoSPARC. (**G**) Angular distribution of particles used for the final 3D reconstruction.

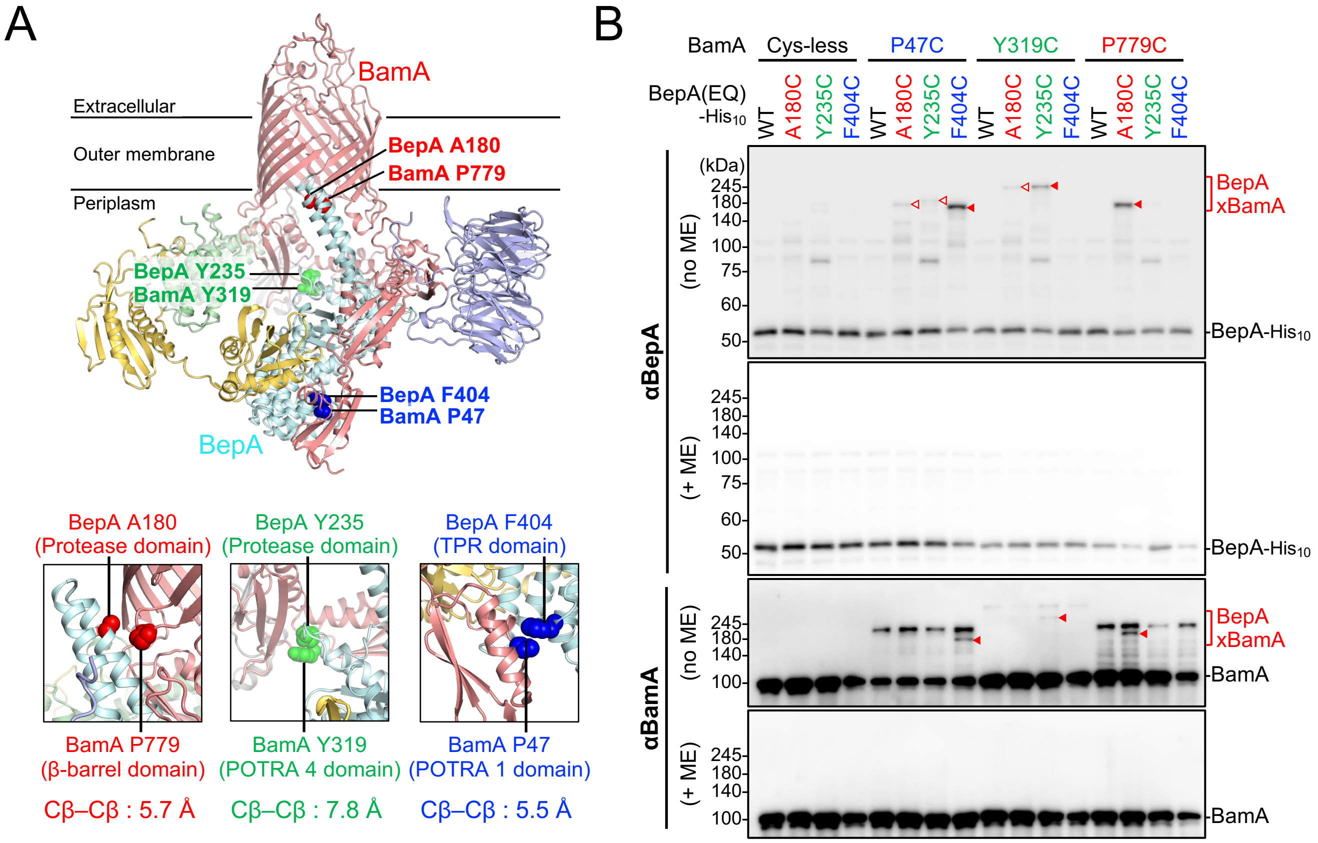


**fig. S10. Disulfide crosslinking between BepA and BamA.**

(**A**) Cysteine-substitution sites in BepA and BamA. In the AlphaFold2 model of the BAM–BepA complex, BamA, BamB, BamC, BamD, BamE, and BepA are in red, light blue, yellow, light green, gray, and pale cyan, respectively. The introduced cysteine positions are shown as red, blue, and green spheres. (**B**) Disulfide crosslinking between BepA and BamA. Cells of RM4748 (Δ*bepA*, $P_{ara}$-*bamA*) carrying a combination of plasmids encoding WT or a Cys-introduced mutant of BepA(E137Q)-$His_{10}$ and BamA as indicated were grown in LB-medium and induced with 1 mM IPTG for 3 h to express BepA(E137Q, Cys)-$His_{10}$ and BamA(Cys). Total cellular proteins were acid-precipitated, solubilized with SDS sample buffer containing NEM (for blocking free thiol groups), treated with or without 2-mercaptoethanol (ME) and analyzed by 7.5% Laemmli SDS-PAGE and immunoblotting with the indicated antibodies.

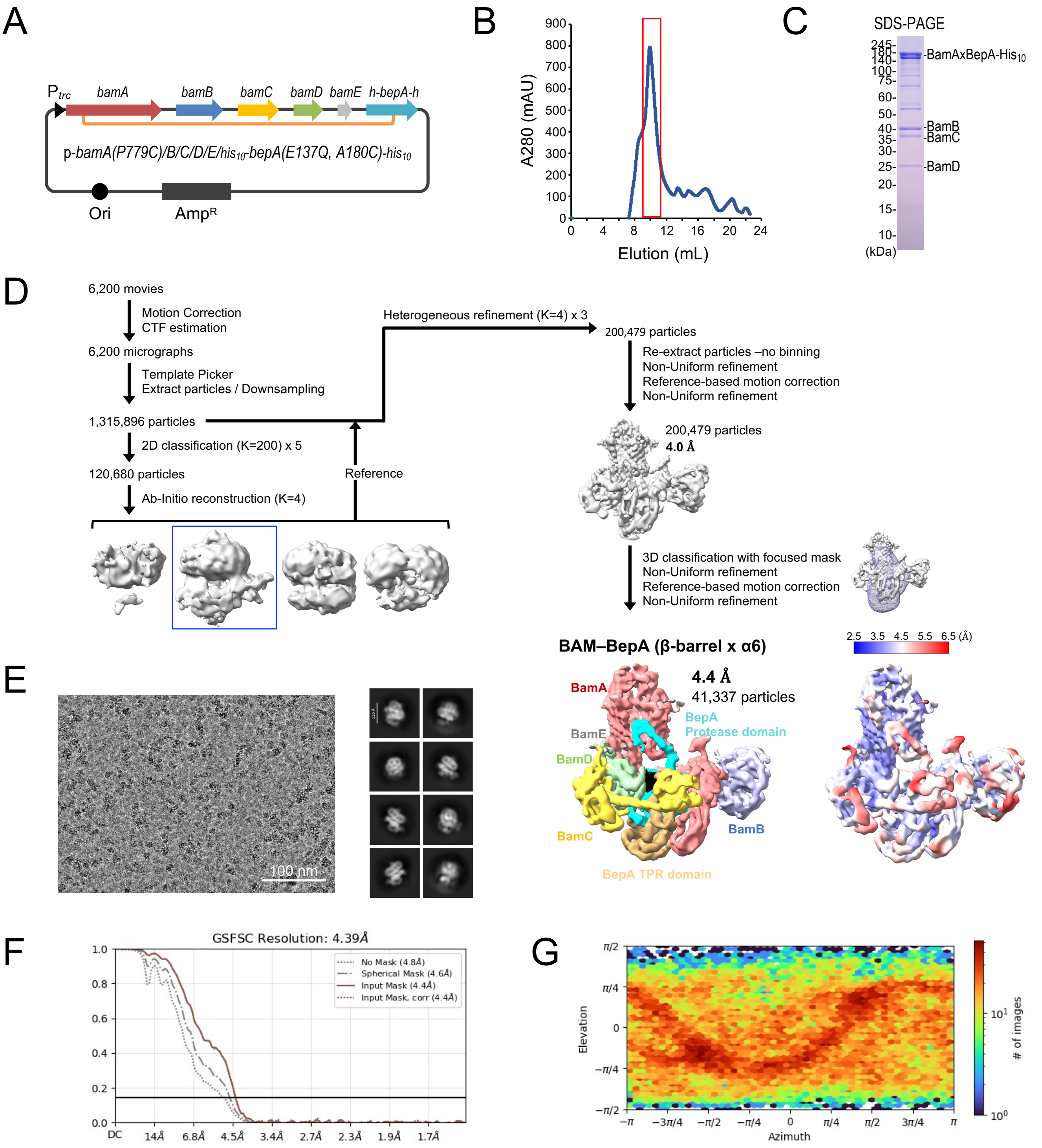


**fig. S11. Cryo-EM analysis of the β-barrel-α6-closslinked BAM–BepA-His complex.**
(**A**) Schematic of a plasmid encoding *bamA(P779C)/B/C/D/E/his$_{10}$-bepA(E137Q, A180C)-his$_{10}$*. N-terminal His$_{10}$-tag is inserted after the signal sequence of BepA. (**B**) Size-exclusion chromatogram of the purified β-barrel-α6-closslinked BAM–BepA-His complex using a Superdex 200 Increase column. (**C**) SDS–PAGE analysis of the purified β-barrel-α6-closslinked BAM–BepA-His complex stained with CBB. (**D**) Cryo-EM data-processing workflow for the β-barrel-α6-closslinked BAM–BepA-His complex. Final cryo-EM map of the β-barrel-α6-closslinked BAM–BepA-His complex. The map is displayed at contour level 0.028. BepA TPR and protease domains are colored in orange and cyan, respectively. BamA, BamB, BamC, BamD, and BamE are in red, light blue, yellow, light green, and gray, respectively. The local resolution of the cryo-EM map was calculated using CryoSPARC and is shown. (**E**) Representative cryo-EM micrograph and 2D classes. (**F**) Gold-standard Fourier shell correlation (GSFSC) curve used for global-resolution estimates within CryoSPARC. (**G**) Angular distribution of particles used for the final 3D reconstruction.

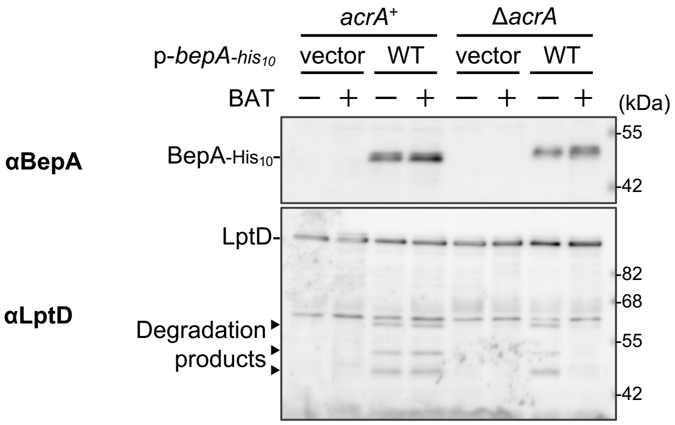


**fig. S12. Batimastat inhibits the LptD degradation activity of BepA.**

Cells of SN56 (Δ*bepA*) or AD2892 (Δ*bepA*, Δ*acrA*) carrying pTWV228-*lptD-his*$_{10}$ and either pHSG575 or pHSG575-*bepA-his*$_{10}$ plasmids were grown, induced as in Supplementary Fig. 1C, and treated with or without 250 μM batimastat. Total cellular proteins were acid-precipitated and analyzed by 7.5 or 10% Laemmli SDS-PAGE and immunoblotting with the indicated antibodies. In the Δ*acrA* background, BAT treatment partially reduced self-cleavage of the C-terminal His$_{10}$-tag of BepA, as indicated by an increased amount of the slower-migrating, uncleaved BepA-His$_{10}$ species.

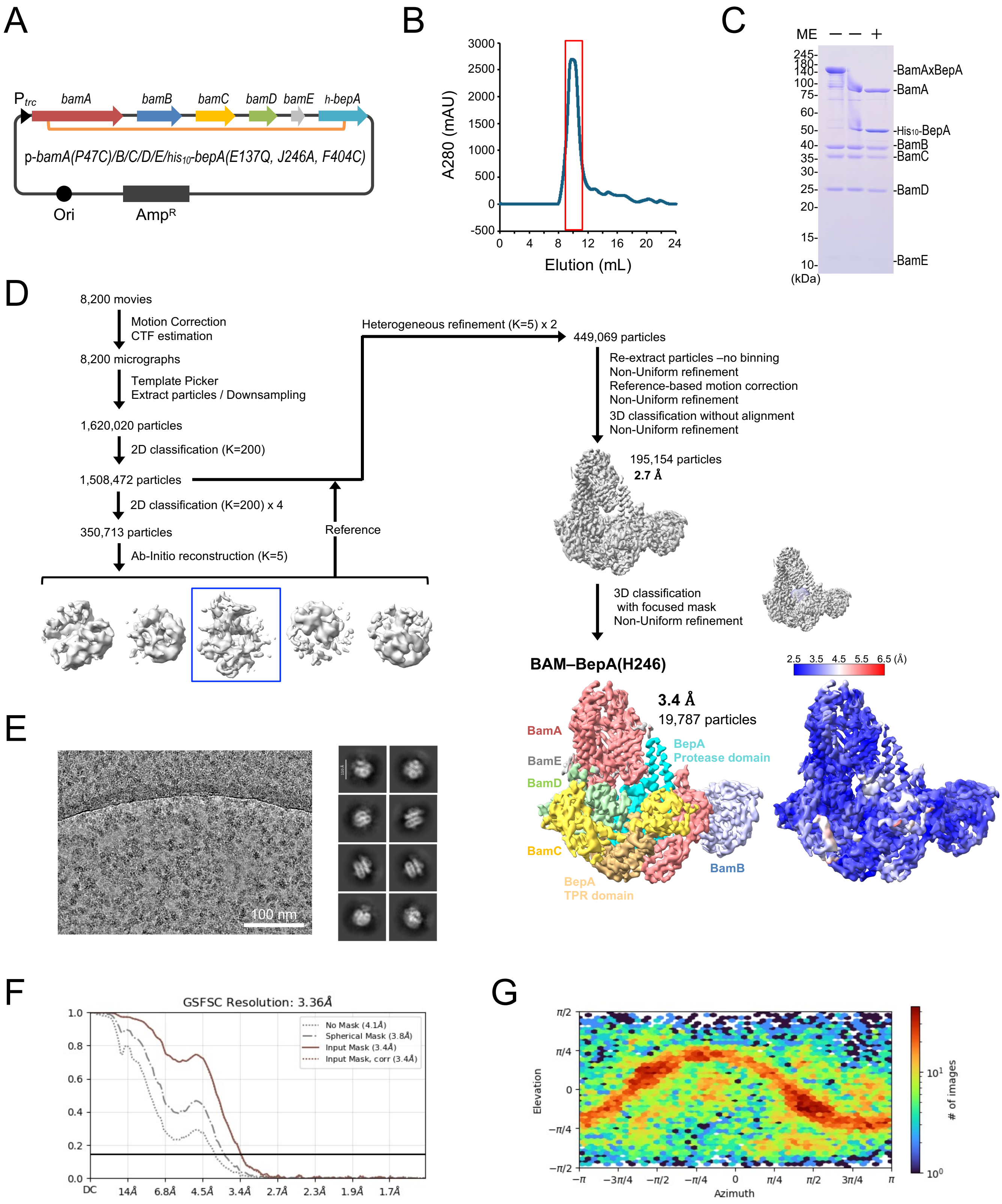


**fig. S13. Cryo-EM analysis of the BAM–BepA(H246A) complex in the presence of Batimastat.**
(**A**) Schematic of a plasmid encoding *bamA(P47C)/B/C/D/E/his₁₀-bepA(E137Q, H246A, F404C)*. N-terminal His$_{10}$-tag is inserted after the signal sequence of BepA. (**B**) Size-exclusion chromatogram of the purified BAM–BepA complex using a Superdex 200 Increase column. (**C**) SDS–PAGE analysis of the purified BAM–BepA complex stained with CBB. (**D**) Cryo-EM data-processing workflow for the BAM–BepA complex. Final cryo-EM maps the BAM–BepA(H246A) in the presence of BAT. The map is displayed at contour level 0.031. BepA TPR and protease domains are colored in orange and cyan, respectively. BamA, BamB, BamC, BamD, and BamE are in red, light blue, yellow, light green, and gray, respectively. The local resolution of the cryo-EM map is calculated using CryoSPARC and is shown. (**E**) Representative cryo-EM micrograph and 2D classes. (**F**) Gold-standard Fourier shell correlation (GSFSC) curve used for global-resolution estimates within CryoSPARC. (**G**) Angular distribution of particles used for the final 3D reconstruction.

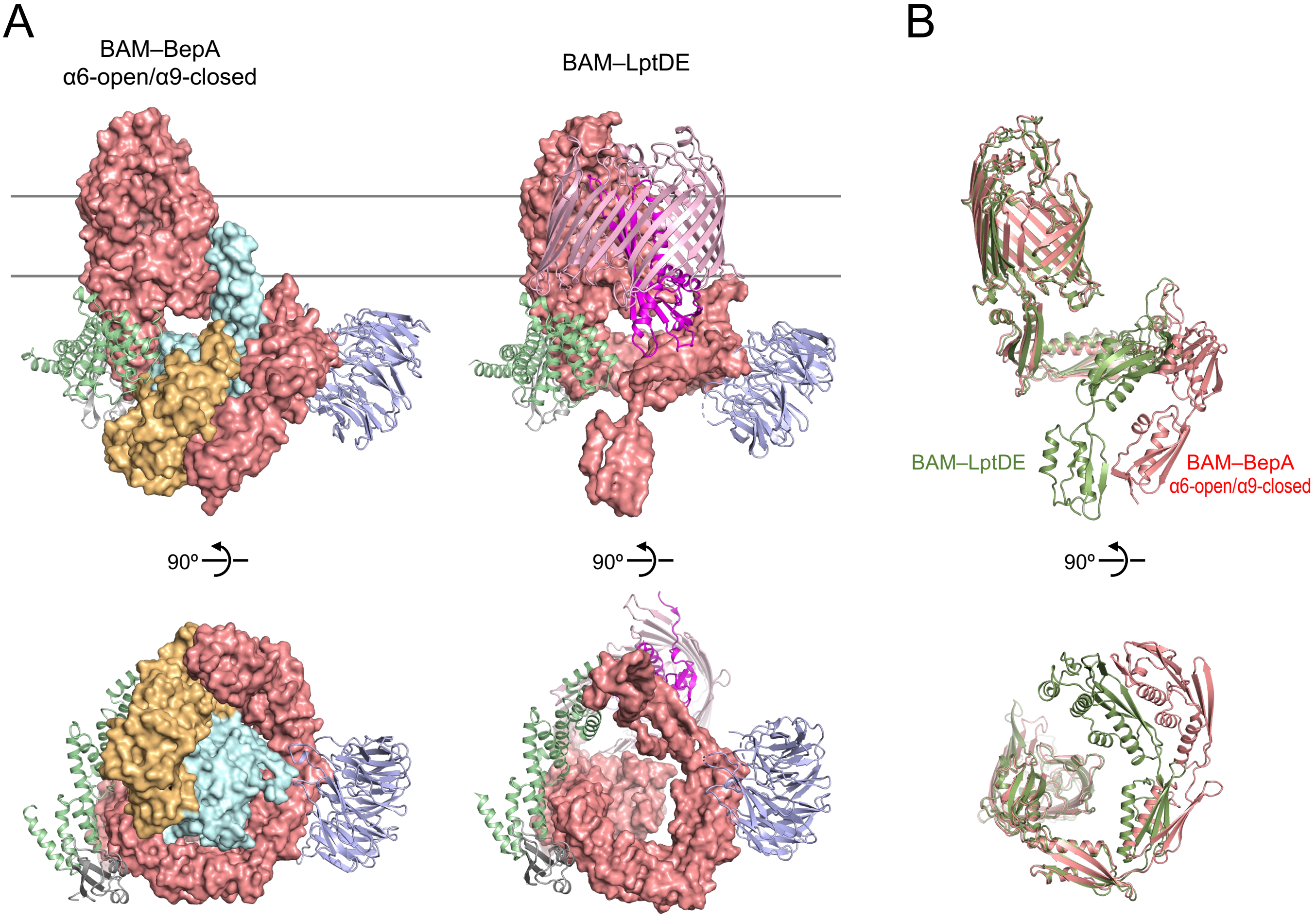


**fig. S14. Comparison of the BAM–BepA α6-open/α9-closed and the BAM-LptDE structures.**
(**A**) Cryo-EM structures of the BAM–BepA and BAM–LptDE complexes. The BAM–BepA α6-open/α9-closed structure was determined in this study (PDB: 27KT), whereas the BAM–LptDE structure was reported previously (PDB: 9P1U). BamA, BamB, BamD, and BamE are shown in red, light blue, light green, and gray, respectively. The BepA protease and TPR domains are shown as pale cyan and orange surfaces, respectively. LptD and LptE are shown as pink and magenta cartoons, respectively. BamC is omitted for clarity. (**B**) Structural comparison of BamA in the BAM–BepA and BAM–LptDE structures. BamA in the BAM–BepA and BAM–LptDE structures is shown in red and green, respectively.

**Table S1. Strains used in this study.**

| Strain | Genotype | Reference |
|---|---|---|
| AD16 | Δ*pro-lac thi*/F' *lacI*$^{q}$ *Z*Δ*M15 Y*$^{+}$ *pro*$^{+}$ | Kihara *et al.* [60] |
| SN56 | AD16, Δ*bepA*::*FRT* | Narita *et al.* [24] |
| AD2892 | AD16, Δ*bepA*::*FRT* Δ*acrA*::*kan* | This study |
| MC4100 | F$^{-}$ *araD139* Δ*(argF-lac)U169 rpsL150 relA1 flbB5301 deoC1 ptsF25 rbsR* | Silhavy *et al.* [61] |
| CU141 | MC4100/F' *lacI*$^{q}$ *lacZ*$^{+}$, *Y*$^{+}$, *A*$^{+}$ | Akiyama *et al.* [62] |
| HM1742 | CU141, *ara*$^{+}$ | Mori and Ito. [44] |
| RM4676 | HM1742,Δ*bepA*::FRT *purC80*::Tn*10* | This study |
| RM4728 | HM1742, *purC80*::Tn*10 kan araC-*$P_{araBAD}$*-bamA* | Miyazaki *et al.* [13] |
| RM4748 | HM1742, Δ*bepA*::*FRT purC80*::Tn*10 kan araC-*$P_{araBAD}$*-bamA* | This study |
| BL21(DE3) | F$^{-}$, *ompT, hsdS*$_{B}$(r$_{B}^{-}$ m$_{B}^{-}$), *gal*(*λc*I 857, *ind*1, *Sam*7, *nin*5, *lacUV*5-T7*gene*1), *dcm*(DE3) | Novagen |

**Table S2. Plasmids used in this study.**

| Plasmid | Vector | Encoded gene and description | Reference or source |
|---|---|---|---|
| pEVOL-pBpF | | p15A-derivative encoding an evolved *M. jannaschii* aminoacyl-tRNA synthetase/suppressor tRNA pair for incorporation of *p*BPA; $Cm^R$ | Young *et al.* [56] |
| pJH113 | | Expression vector; $P_{tac}$, $Amp^R$ | Roman-Hernandez *et al.* [63] |
| pRM1261 | pJH113 | *bamA(C690S, C700S)/bamB/bamC/bamD/bamE-his*$_8$ | Miyazaki *et al.* [13] |
| pRM1596 | pJH113 | *bamA(P47C, C690S, C700S)/bamB/bamC/bamD/bamE-his*$_8$ | This study |
| pRM1598 | pJH113 | *bamA(C690S, C700S, P779C)/bamB/bamC/bamD/bamE-his*$_8$ | This study |
| pRM1752 | pJH113 | *bamA(C690S, C700S, P779C)/bamB/bamC/bamD/bamE/his*$_{10}$*-bepA(E137Q, A180C)-his*$_{10}$ | This study |
| pRM1763 | pJH113 | *bamA(P47C, C690S, C700S)/bamB/bamC/bamD/bamE/his*$_{10}$*-bepA(E137Q, F404C)-his*$_{10}$ | This study |
| pRM1949 | pJH113 | *bamA(P47C, C690S, C700S)/bamB/bamC/bamD/bamE/his*$_{10}$*-bepA(E137Q, F404C)* | This study |
| pRM2111 | pJH113 | *bamA(P47C, C690S, C700S)/bamB/bamC/bamD/bamE/his*$_{10}$*-bepA(E137Q, H246A, F404C)* | This study |
| pUC18 | | Expression vector; $P_{lac}$, $Amp^R$ | Takara Bio |
| pUC-bepA(E137Q)-his$_{10}$ | pUC18 | *bepA(E137Q)-his*$_{10}$ | Narita *et al.* [24] |
| pRM1091 | pUC18 | *bepA(E137Q, Q51amb)-his*$_{10}$ | This study |
| pRM1094 | pUC18 | *bepA(E137Q, S63amb)-his*$_{10}$ | This study |
| pRM1098 | pUC18 | *bepA(E137Q, P93amb)-his*$_{10}$ | This study |
| pRM1120 | pUC18 | *bepA(E137Q, R148amb)-his*$_{10}$ | This study |
| pRM1102 | pUC18 | *bepA(E137Q, V162amb)-his*$_{10}$ | This study |
| pRM1104 | pUC18 | *bepA(E137Q, I168amb)-his*$_{10}$ | This study |
| pRM1106 | pUC18 | *bepA(E137Q, Q176amb)-his*$_{10}$ | This study |
| pRM1123 | pUC18 | *bepA(E137Q, A180amb)-his*$_{10}$ | This study |
| pRM1125 | pUC18 | *bepA(E137Q, T185amb)-his*$_{10}$ | This study |
| pRM1127 | pUC18 | *bepA(E137Q, R190amb)-his*$_{10}$ | This study |
| pRM1107 | pUC18 | *bepA(E137Q, M193amb)-his*$_{10}$ | This study |
| pRM1108 | pUC18 | *bepA(E137Q, Q198amb)-his*$_{10}$ | This study |
| pRM1130 | pUC18 | *bepA(E137Q, R214amb)-his*$_{10}$ | This study |
| pRM1131 | pUC18 | *bepA(E137Q, E227amb)-his*$_{10}$ | This study |
| pRM1133 | pUC18 | *bepA(E137Q, Y235amb)-his*$_{10}$ | This study |

| | | | |
|---|---|---|---|
| pRM1111 | pUC18 | *bepA(E137Q, E241amb)-his*$_{10}$ | This study |
| pRM1113 | pUC18 | *bepA(E137Q, S251amb)-his*$_{10}$ | This study |
| pRM1114 | pUC18 | *bepA(E137Q, D255amb)-his*$_{10}$ | This study |
| pRM1115 | pUC18 | *bepA(E137Q, N258amb)-his*$_{10}$ | This study |
| pUC-bepA(E137Q/F404amb)-his$_{10}$ | pUC18 | *bepA(E137Q, F404amb)-his*$_{10}$ | Daimon *et al.* [30] |
| pUC-bepA-his$_{10}$ | pUC18 | *bepA-his*$_{10}$ | Narita *et al.* [24] |
| pRM1511 | pUC18 | *bepA(Q51amb)-his*$_{10}$ | This study |
| pRM1496 | pUC18 | *bepA(S63amb)-his*$_{10}$ | This study |
| pRM1497 | pUC18 | *bepA(R148amb)-his*$_{10}$ | This study |
| pRM1498 | pUC18 | *bepA(V162amb)-his*$_{10}$ | This study |
| pRM1499 | pUC18 | *bepA(I168amb)-his*$_{10}$ | This study |
| pRM1500 | pUC18 | *bepA(Q176amb)-his*$_{10}$ | This study |
| pRM1501 | pUC18 | *bepA(A180amb)-his*$_{10}$ | This study |
| pRM1502 | pUC18 | *bepA(R190amb)-his*$_{10}$ | This study |
| pRM1503 | pUC18 | *bepA(M193amb)-his*$_{10}$ | This study |
| pRM1504 | pUC18 | *bepA(R214amb)-his*$_{10}$ | This study |
| pRM1505 | pUC18 | *bepA(E227amb)-his*$_{10}$ | This study |
| pRM1506 | pUC18 | *bepA(Y235amb)-his*$_{10}$ | This study |
| pRM1507 | pUC18 | *bepA(E241amb)-his*$_{10}$ | This study |
| pRM1508 | pUC18 | *bepA(S251amb)-his*$_{10}$ | This study |
| pRM1509 | pUC18 | *bepA(D255amb)-his*$_{10}$ | This study |
| pRM1510 | pUC18 | *bepA(N258amb)-his*$_{10}$ | This study |
| pHSG575 | | Expression vector; $P_{lac}$, $Cm^R$ | NBRP |
| pYD552 | pHSG575 | *bepA-his*$_{10}$ | This study |
| pYD556 | pHSG575 | *bepA(E137Q)-his*$_{10}$ | This study |
| pSTD1554 | pHSG575 | *bepA(N105C)-his*$_{10}$ | This study |
| pSTD1531 | pHSG575 | *bepA(F107C)-his*$_{10}$ | This study |
| pSTD1548 | pHSG575 | *bepA(V115C)-his*$_{10}$ | This study |
| pSTD1505 | pHSG575 | *bepA(S132C)-his*$_{10}$ | This study |
| pSTD1507 | pHSG575 | *bepA(V133C)-his*$_{10}$ | This study |
| pSTD1552 | pHSG575 | *bepA(L165C)-his*$_{10}$ | This study |
| pSTD1550 | pHSG575 | *bepA(I242C)-his*$_{10}$ | This study |
| pSTD1530 | pHSG575 | *bepA(A277C)-his*$_{10}$ | This study |
| pSTD1529 | pHSG575 | *bepA(D298C)-his*$_{10}$ | This study |

| | | | |
|---|---|---|---|
| pRM1474 | pHSG575 | *bepA(Δα6)-his*$_{10}$ | This study |
| pRM1476 | pHSG575 | *bepA(Δα6, E137Q)-his*$_{10}$ | This study |
| pSTD1636 | pHSG575 | *bepA(*L60C, *L170C)-his*$_{10}$ | This study |
| pSTD1640 | pHSG575 | *bepA(*L60C, *L170C, E137Q)-his*$_{10}$ | This study |
| pSTD1519 | pHSG575 | *bepA(H246A)-his*$_{10}$ | This study |
| pSTD1525 | pHSG575 | *bepA(S132C, H246A)-his*$_{10}$ | This study |
| pSTD1527 | pHSG575 | *bepA(V133C, H246A)-his*$_{10}$ | This study |
| pSTD1570 | pHSG575 | *bepA(Δα9)-his*$_{10}$ | This study |
| pSTD1625 | pHSG575 | *bepA(S132C, Δα9)-his*$_{10}$ | This study |
| pSTD1627 | pHSG575 | *bepA(V133C, Δα9)-his*$_{10}$ | This study |
| pTWV228 | | Expression vector; P$_{lac}$, Amp$^{R}$ | Takara Bio |
| pRM265 | pTWV228 | *lptD-his*$_{10}$ (using native SD sequence) | Daimon *et al.* [30] |
| pRM816 | pTWV228 | *lptD-his*$_{10}^{SD}$ (using strong SD sequence) | Miyazaki *et al.* [31] |
| pRM811 | pTWV228 | *lptD(Y331C)-his*$_{10}^{SD}$ | Miyazaki *et al.* [31] |
| pRM812 | pTWV228 | *lptD(E391C)-his*$_{10}^{SD}$ | Miyazaki *et al.* [31] |
| pRM815 | pTWV228 | *lptD(V430C)-his*$_{10}^{SD}$ | Miyazaki *et al.* [31] |
| pSTD689 | | Expression vector; P$_{lac}$, Spc$^{R}$ | Kanehara *et al.* [45] |
| pRM290 | pSTD689 | *bepA* | Daimon *et al.* [30] |
| pRM291 | pSTD689 | *bepA(E137Q)* | Daimon *et al.* [30] |
| pRM786 | pSTD689 | *bepA(E137Q, N105C)* | Miyazaki *et al.* [31] |
| pRM1568 | pSTD689 | *bepA(I168C)* | This study |
| pRM1569 | pSTD689 | *bepA(R190C)* | This study |
| pRM1572 | pSTD689 | *bepA(E137Q, I168C)* | This study |
| pRM1573 | pSTD689 | *bepA(E137Q, R190C)* | This study |
| pRM866 | pSTD689 | *bepA-his*$_{10}$ | This study |
| pRM868 | pSTD689 | *bepA(E137Q)-his*$_{10}$ | This study |
| pRM1350 | pSTD689 | *bepA(E137Q, A180C)-his*$_{10}$ | This study |
| pRM1351 | pSTD689 | *bepA(E137Q, Y235C)-his*$_{10}$ | This study |
| pRM888 | pSTD689 | *bepA(E137Q, F404C)-his*$_{10}$ | This study |
| pRM1370 | pSTD689 | *bepA(E137Q, A180C)-his*$_{10}$ | This study |
| pRM1369 | pSTD689 | *bepA(E137Q, Y235C)-his*$_{10}$ | This study |
| pRM901 | pSTD689 | *bepA(E137Q, F404C)-his*$_{10}$ | This study |
| pRM1612 | pSTD689 | *his*$_{10}$*-GGSG-bepA(E137Q)-his*$_{10}$ | This study |
| pRM1615 | pSTD689 | *his*$_{10}$*-GGSG-bepA(E137Q, A180C)-his*$_{10}$ | This study |
| pRM1619 | pSTD689 | *his*$_{10}$*-GGSG-bepA(E137Q, F404C)-his*$_{10}$ | This study |
| pUC118 | | Expression vector; P$_{lac}$, Amp$^{R}$ | Takara Bio |

| | | | |
|---|---|---|---|
| pRM823 | pUC118 | *bamA* | Miyazaki *et al.* [31] |
| pRM864 | pUC118 | *bamA(C690S, C700S)* | Miyazaki *et al.* [13] |
| pRM1333 | pUC118 | *bamA(P47C, C690S, C700S)* | This study |
| pRM1341 | pUC118 | *bamA(Y319C, C690S, C700S)* | This study |
| pRM1345 | pUC118 | *bamA(P779C, C690S, C700S)* | This study |

**Supplementary Table 3. Cryo-EM data collection and refinement statistics.**

| dataset | BAM–BepA in ND | BAM–BepA in DDM | | BAM–BepA (H246A) in DDM |
|---|---|---|---|---|
| Structure | α6/α9-closed | α6/α9-closed | α6-open /α9-closed | α6/α9-open |
| | (EMD-81243) | (EMD-81235) | (EMD-81228) | (EMD-82637) |
| | (PDB ID: 27LP) | (PDB ID: 27LF) | (PDB ID: 27KT) | (PDB ID: 44IO) |
| **Data collection and processing** | | | | |
| Microscope | CRYO ARM 300 | Glacios | Glacios | CRYO ARM 300 |
| Magnification | x60,000 | x190,000 | x190,000 | x60,000 |
| Voltage (kV) | 300 | 200 | 200 | 300 |
| Electron exposure (e–/Å$^2$) | 49.98 | 50 | 50 | 50 |
| Defocus range (μm) | -1.4 to -1.6 | -0.8 to -1.4 | -0.8 to -1.4 | -1.4 to -1.6 |
| Pixel size (Å) | 0.752 | 0.74 | 0.74 | 0.752 |
| Symmetry imposed | C1 | C1 | C1 | C1 |
| Initial particle images (no.) | 2,741,395 | 1,247,906 | 1,247,906 | 1,623,020 |
| Final particle images (no.) | 13,551 | 49,087 | 22,410 | 19,787 |
| Map resolution (Å) | 3.88 | 4.11 | 3.89 | 3.36 |
| FSC threshold | 0.143 | 0.143 | 0.143 | 0.143 |
| Map resolution range (Å) | 48.43 to 1.89 | 46.87 to 2.28 | 50.52 to 2.34 | 41.80 to 1.90 |
| **Refinement** | | | | |
| Initial model used (PDB code) | 25FQ AlphaFold model | 25FQ AlphaFold model | 25FQ AlphaFold model | 25FQ AlphaFold model |
| Model composition | | | | |
| Non-hydrogen atoms | 15,137 | 17,035 | 17,356 | 17,314 |
| Protein residues | 1,937 | 2185 | 2,232 | 2,225 |
| Ligands | 1 | 1 | 1 | 1 |
| *B* factors (Å$^2$) | | | | |
| Protein | 143.54 | 174.29 | 123.31 | 121.56 |
| Ligand | 200.06 | 309.78 | 132.54 | 200.06 |
| R.m.s. deviations | | | | |
| Bond lengths (Å) | 0.007 | 0.008 | 0.005 | 0.006 |
| Bond angles (°) | 1.193 | 1.164 | 1.005 | 1.012 |
| **Validation** | | | | |
| MolProbity score | 2.28 | 1.86 | 1.78 | 1.57 |
| Clashscore | 17.32 | 15.99 | 12.80 | 10.20 |
| Poor rotamer (%) | 2.79 | 1.04 | 1.57 | 1.14 |
| Ramachandran plot | | | | |
| Favored (%) | 96.72 | 97.23 | 97.93 | 98.15 |
| Allowed (%) | 3.28 | 2.77 | 2.07 | 1.81 |
| Disallowed (%) | 0 | 0 | 0 | 0.05 |

**Supplementary Table 3. Cryo-EM data collection and refinement statistics. (Continued)**

| dataset | BAM–BepA-His in ND | BAM–BepA-His in DDM | BAM–BepA-His (β-barrel x α6) in DDM |
|---|---|---|---|
| Structure | α6/α9-closed | α6-open /α9-closed | α6-open /α9-closed |
| | (EMD-81262) | (EMD-81260) | (EMD-81261) |
| | (PDB ID: 27ML) | (PDB ID: 27MJ) | (PDB ID: 27MK) |
| **Data collection and processing** | | | |
| Microscope | Titan Krios | CRYO ARM 300 | CRYO ARM 300 |
| Magnification | x165,000 | x60,000 | x60,000 |
| Voltage (kV) | 300 | 300 | 300 |
| Electron exposure (e−/Å$^2$) | 50 | 49.92 | 50.88 |
| Defocus range (μm) | -0.6 to -1.6 | -1.4 to -1.6 | -1.4 to -1.6 |
| Pixel size (Å) | 0.76 | 0.752 | 0.752 |
| Symmetry imposed | C1 | C1 | C1 |
| Initial particle images (no.) | 1,452,797 | 1,577,906 | 1,315,897 |
| Final particle images (no.) | 9,882 | 8,952 | 41,337 |
| Map resolution (Å) | 4.12 | 4.04 | 4.39 |
| FSC threshold | 0.143 | 0.143 | 0.143 |
| Map resolution range (Å) | 52.15 to 2.48 | 51.58 to 2.24 | 57.16 to 2.92 |
| **Refinement** | | | |
| Initial model used (PDB code) | 25FQ<br>AlphaFold model | 25FQ<br>AlphaFold model | 25FQ<br>AlphaFold model |
| Model composition | | | |
| Non-hydrogen atoms | 14,993 | 17,340 | 17,465 |
| Protein residues | 1,918 | 2,230 | 2,241 |
| Ligands | 1 | 1 | 1 |
| *B* factors (Å$^2$) | | | |
| Protein | 144.54 | 145.31 | 241.94 |
| Ligand | 256.80 | 149.83 | 277.54 |
| R.m.s. deviations | | | |
| Bond lengths (Å) | 0.006 | 0.005 | 0.005 |
| Bond angles (°) | 1.121 | 1.053 | 1.039 |
| **Validation** | | | |
| MolProbity score | 1.71 | 1.70 | 1.63 |
| Clashscore | 11.34 | 14.92 | 12.88 |
| Poor rotamer (%) | 0.56 | 0.16 | 0.21 |
| Ramachandran plot | | | |
| Favored (%) | 97.26 | 97.92 | 97.98 |
| Allowed (%) | 2.74 | 2.08 | 2.02 |
| Disallowed (%) | 0 | 0 | 0 |
